# Organization, genomic targeting and assembly of three distinct SWI/SNF chromatin remodeling complexes in *Arabidopsis*

**DOI:** 10.1101/2022.11.24.517835

**Authors:** Wei Fu, Yaoguang Yu, Jie Shu, Zewang Yu, Tao Zhu, Yixiong Zhong, Zhihao Zhang, Zhenwei Liang, Yuhai Cui, Chen Chen, Chenlong Li

## Abstract

Switch defective/sucrose non-fermentable (SWI/SNF) complexes are evolutionarily conserved multi-subunit machines that play vital roles in chromatin architecture regulation for modulating gene expression via sliding or ejection of nucleosomes in eukaryotes. In plants, perturbations of SWI/SNF subunits often result in severe developmental disorders. However, the subunit composition, pathways of assembly, and genomic targeting of the plant SWI/SNF complexes remain undefined. Here, we reveal the organization, genomic targeting and assembly of three distinct Arabidopsis SWI/SNF complexes: <u>B</u>RAHMA-<u>A</u>ssociated <u>S</u>WI/SNF complexes (BAS), <u>S</u>PLAYED-<u>A</u>ssociated <u>S</u>WI/SNF complexes (SAS) and <u>M</u>INUSCULE-<u>A</u>ssociated <u>S</u>WI/SNF complexes (MAS). We show that BAS complexes are equivalent to human ncBAF, whereas SAS and MAS complexes evolve in multiple subunits unique to plants, suggesting a plant-specific functional evolution of SWI/SNF complexes. We further demonstrate overlapping and specific genomic targeting of the three plant SWI/SNF complexes on chromatin and reveal that SAS complexes are necessary for the correct genomic localization of the BAS complexes. Finally, we define the role of core module subunit in the assembly of the plant SWI/SNF complexes and highlight that ATPase module subunit is required for global complex stability and the interaction of core module subunits in SAS and BAS complexes in Arabidopsis. Together, our work highlights the divergence of SWI/SNF chromatin remodelers during the eukaryote evolution and provides a comprehensive landscape for understanding the plant SWI/SNF complexes organization, assembly, genomic targeting, and function.

**One-sentence summary:** Comprehensively define the organization, genomic targeting and assembly of three distinct SWI/SNF chromatin remodeling complexes in Arabidopsis

## Introduction

Switch defective/sucrose non-fermentable (SWI/SNF) complexes are chromatin remodelers that play essential roles in modulating chromatin architecture to enable DNA accessibility and gene expression in an ATP-hydrolysis-dependent manner (Clapier and Cairns, 2009; Ho and Crabtree, 2010; Hargreaves and Crabtree, 2011; Ryan and Owen-Hughes, 2011; Clapier et al., 2017). These complexes are multi-subunit machineries, including one ATPase catalytic subunit and multiple additional regulatory core subunits, evolutionally conserved among yeasts, animals and plants (Ho and Crabtree, 2010; Sarnowska et al., 2016). In *Saccharomyces cerevisiae*, there are two sub-families of SWI/SNF remodelers, Swi/Snf and RSC. The yeast Swi/Snf complex consists of 12 proteins, which was the first chromatin remodeler discovered (Smith et al., 2003; Dutta et al., 2017), whereas the RSC complex is composed of 16 proteins (Wagner et al., 2020). The two sub-complexes contain three common subunits, ARP7/9 and Rtt102, and the other subunits are specialized in the two sub-complexes (Dutta et al., 2017; Wagner et al., 2020). In humans, the subunits of SWI/SNF complexes are combinatorially assembled into three classes of complexes: canonical BAF (cBAF), polybromo-associated BAF (PBAF), and non-canonical BAF (ncBAF) (Gatchalian et al., 2018; Mashtalir et al., 2018). These three SWI/SNF sub-complexes share a set of core subunits, such as SMARCA2/SMARCA4, SMARCC1/2, SMARCD, ACTL6A/B, and BCL7A/B/C, but are distinguished by the inclusion of subtype-specific ones: i.e., DPF1, DPF2, ARID1A, and ARID1B for cBAF; PBRM1, PHF10, ARID2, and BRD7 for PBAF; and BRD9 and GLTSCR1/1L for ncBAF (Gatchalian et al., 2018; Mashtalir et al., 2018).

In the flowering plant *Arabidopsis thaliana*, through genetic and molecular studies, a number of SWI/SNF subunits have been identified, including four ATPases (BRAHMA (BRM), SPLAYED (SYD), MINUSCULE 1 (MINU1), and MINU2; four SWI3 subunits (SWI3A-SWI3D); two SWI/SNF associated proteins 73 (SWP73A and SWP73B); two actin-related proteins (ARP4 and ARP7); a single SNF5 subunit called BUSHY (BSH); an ARID paralog LEAF AND FLOWER RELATED (LFR) (Wagner D, 2002; Farrona et al., 2004; Hurtado et al., 2006; Mlynarova et al., 2007; Wang et al., 2009; Sang et al., 2012). In addition, we recently biochemically isolated novel subunits of the BRM-containing SWI/SNF complexes in Arabidopsis, including two BRM-interacting proteins (BRIP1 and BRIP2) and three bromodomain-containing proteins, BRD1, BRD12, and BRD13 (Yu et al., 2020; Yu et al., 2021). More recently, MINU-containing SWI/SNF complexes were also reported (Diego-Martin et al., 2022). Plant SWI/SNF subunits serve critical roles in cell differentiation, development, and response to various environmental signals, and their mutations result in severe defects in leaf development, root stem cell maintenance, flower patterning and timing, embryo development, and vegetative to adult phase transition (Sarnowski et al., 2005; Ho and Crabtree, 2010; Li et al., 2015; Wu et al., 2015; Yang et al., 2015; Zhao et al., 2015; Sarnowska et al., 2016; Xu et al., 2016). However, owing in large part to limitations in understanding plant SWI/SNF subunit composition, the mechanisms by which these mutations alter plant SWI/SNF complexes function on chromatin and subsequently lead to impaired development and responses to environmental stimulus remain unknown. Particularly, how many different types of SWI/SNF sub-complexes are presented in plants and the subunit composition of each sub-complexes is unclear. Finally, how the activities and genomic targeting of different SWI/SNF subtypes are coordinated with each other to regulate chromatin structure and gene expression is also poorly understood.

Another significant barrier to our mechanistic understanding of the functions of SWI/SNF complexes lies in the need for more information regarding the role of each subunit in complex assembly. Recent studies have begun to reveal the assembly steps of the subunits of SWI/SNF complexes. It was reported that mammalian SWI/SNF complex assembly is triggered by the formation of the initial BAF core that is composed of two SMARCC and one SMARCD subunits. This initial core then acts as a platform for independent docking of the subcomplex-specific subunits to form the core module. Finally, the core module recruits the ATPase SMARCA2/4 to finalize complex assembly (Gatchalian et al., 2018; Mashtalir et al., 2018). In line with this model, the removal of core module subunits SMAECCs resulted in near-complete degradation of all three SWI/SNF complexes components, whereas removal of the ATPase module does not disrupt the formation of the core module (Mashtalir et al., 2018; Michel et al., 2018; Pan et al., 2019). Thus, the core module is required for assembly toward fully formed SWI/SNF complexes in mammalian, but the ATPase module is the last to be incorporated into SWI/SNF complexes and, therefore, not necessary for the core module assembly in mammals (Mashtalir et al., 2018; Michel et al., 2018; Pan et al., 2019). In *Arabidopsis*, our recent studies showed that BRIP1/2 and BRD1/2/13, the homologs of human ncBAF core module subunits GLTSCR1/1L and BRD9, are required for the assembly of SWI/SNF complexes (Yu et al., 2020). However, whether plant ATPases use the same strategy as their mammalian counterparts or a different one for complex assembly and thus the underlying mechanisms are still unknown.

In this study, we examined the organization, genomic targeting, and assembly of the three non-redundant final form Arabidopsis SWI/SNF complexes: <u>B</u>RAHMA-<u>A</u>ssociated <u>S</u>WI/SNF complexes (BAS), <u>S</u>PLAYED-<u>A</u>ssociated <u>S</u>WI/SNF complexes (SAS) and <u>M</u>INUSCULE-<u>A</u>ssociated <u>S</u>WI/SNF complexes (MAS). BAS complexes are equivalent to human ncBAF, whereas SAS and MAS complexes evolve in multiple subunits unique to plants, suggesting a plant-specific functional evolution of SWI/SNF complexes. We further demonstrate both overlapping and specific genomic targeting of the three plant SWI/SNF complexes on chromatin and reveal a requirement of SAS complexes for the correct genomic localization of the BAS complexes. Finally, by focusing on the SAS and BAS complexes, we define the role of core module subunit in the assembly of the plant SWI/SNF complexes and unexpectedly establish that ATPase module subunit (SYD and BRM) is required for the stability and interaction of core module subunits in SAS and BAS complexes in Arabidopsis. Together, these studies highlight the divergence of SWI/SNF chromatin remodelers during the eukaryote evolution and lay the groundwork for comprehensive understanding the plant SWI/SNF complexes organization, assembly, genomic targeting, and function.

## Results

### Arabidopsis has three biochemically distinct SWI/SNF sub-complexes

To comprehensively define the potential plant SWI/SNF sub-complexes, their subunits organization and assembly, we started by performing immuno-purification followed by mass spectrometry analysis (IP-MS) using stable transgenic *Arabidopsis* lines that expressed a green fluorescent protein (GFP)-tagged ATPase subunit, BRM (Li et al., 2016), SYD (Shu et al., 2021) or MINU2 (this study), driven by its native promoter in corresponding null mutant background. This analysis revealed that the three ATPases were constituted into three distinct sub-assemblies (Figure 1 A-B, and Supplemental Figure S1, A-C). Specifically, we found that the BRM-containing complexes uniquely lack core, evolutionarily conserved ARID paralog, LFR, incorporate selective paralogs (that is, SWI3C but not SWI3A/B/D, and SWP73A) and contain a set of complex-specific subunits that are not shared by SYD- or MINU-complex, the GLTSCR1/1L paralogs (BRIP1/2) and BRD1/2/13. In contrast, the SYD-containing complexes selectively contain paralog SWI3D but not SWI3A/B/C, and include a complex-specific subunit, encoded by *<u>S</u>YD sub-<u>S</u>WI/SNF Interacting <u>P</u>rotein 1/2/3* (*SSIP1*, *SSIP2,* or *SSIP3*). Finally, consistent with the recent report (Diego-Martin et al., 2022), the MINU-containing complexes do not contain the GIF2 subunit (SS18 paralogs) found in both BRM- and SYD-containing complexes, incorporate selective paralogs (that is, SWI3A/B but not SWI3C and SWI3D) and include a number of complex-specific subunits, including BSH, BRD5, SHH2, TPF1/2, OPF1/2, PSA1, and PSA2.

**Figure 1.**
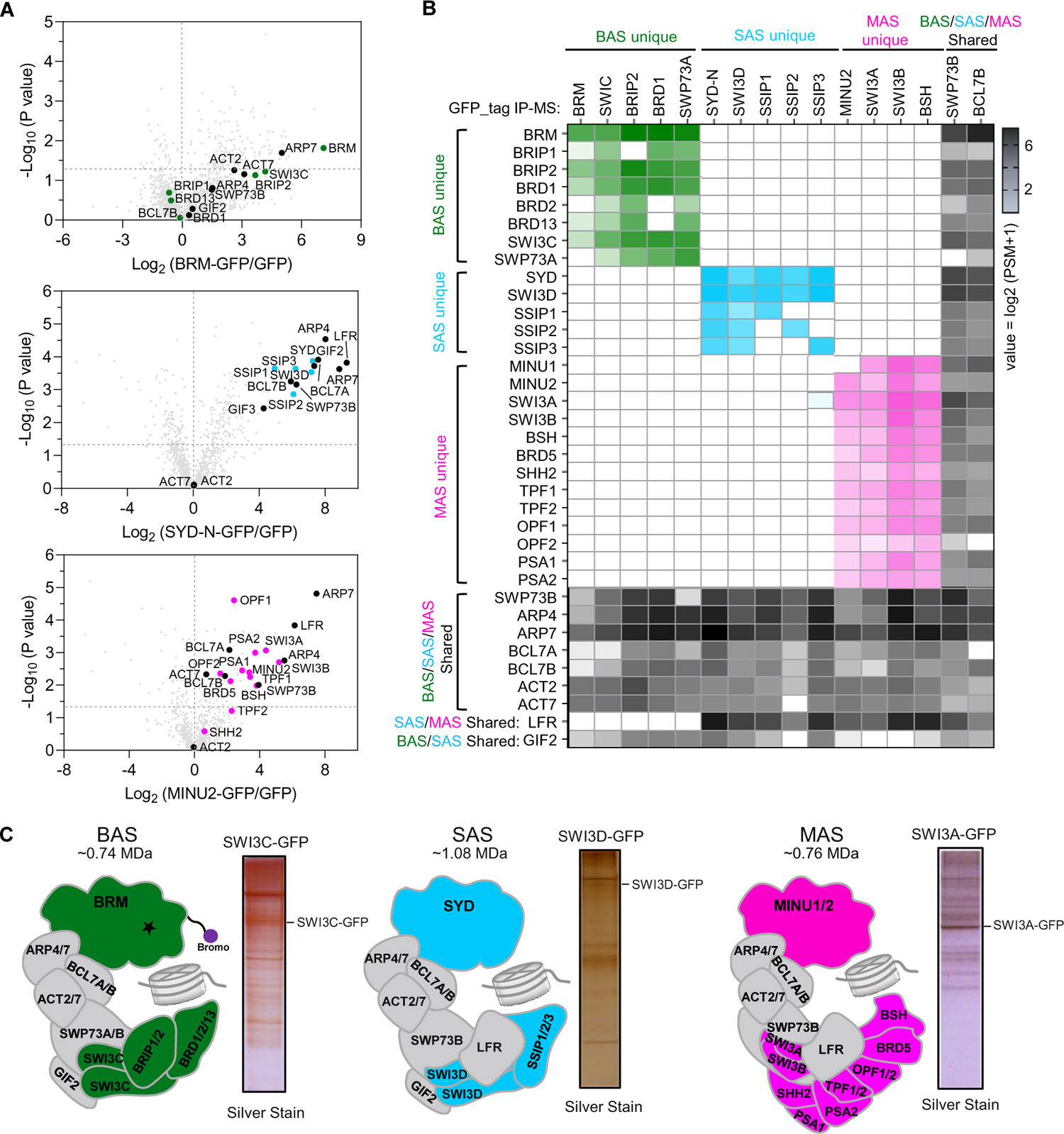
Three distinct SWI/SNF sub-complexes in *Arabidopsis*. **A**, Volcano plots displaying SWI/SNF subunits that are enriched in GFP immunoprecipitations from BRM-GFP, SYD-GFP, MINU2-GFP relative to GFP from two independent experiments. *P* values were calculated by two-tailed Student’s t-test. Subunits of the *Arabidopsis* SWI/SNF complexes are highlighted using green dots (BRM unique subunits), blue dots (SYD unique subunits), magenta dots (MINU unique subunits) and black dots (BRM, SYD and MINU2 shared subunits). **B**, Heatmap showing the mean log_2_(PSM+1) values (from two biological replications) of SWI/SNF complex subunits identified by IP-MS in BRM, SWI3C, BRIP2, BRD1, SWP73A, SYD-N, SWI3D, SSIP1/2/3, MINU2, SWI3A, SWI3B, BSH, SWP73B and BCL7B. **C**, Diagrams showing the organization of BAS, SAS and MAS complexes, respectively. Sliver-stained gels of GFP immunoprecipitations from SWI3C-GFP, SWI3D-GFP and SWI3A-GFP were shown. The specific subunits of each sub-complex were marked with different colors, and the common subunits of different sub-complexes were marked with gray.

To further confirm the three distinct SWI/SNF sub-complexes in Arabidopsis, we generated transgenic lines stably expressing GFP-tagged subunits specific to the BRM-complexes, the SYD-complexes, or the MINU-complexes or subunits shared by the three sub-complexes. We then identified proteins that co-purified with each subunit by mass spectrometry. Silver staining of proteins isolated from transgenic lines stably expressing GFP-tagged SWI3C, SWI3D and SWI3A again demonstrated three distinct SWI/SNF sub-complexes in *Arabidopsis* (Figure 1C and Supplemental Figure S1D). Furthermore, mass-spectrometry showed that SWI3C, BRIP2, BRD1, and SWP73A immunoprecipitation enriched the identical subunits immunoprecipitated by BRM, whereas none of the SYD-specific or MINU-specific subunits were co-immunoprecipitated, indicating that they are unique components to the BRM-containing complexes (Figure 1B and Supplemental Figure S1E). Likewise, the immunoprecipitation of SYD-specific or MINU-specific subunits did not catch subunits unique to the other two subcomplexes (Figure 1B and Supplemental Figure S1E). In contrast, SWP73B and BCL7B, the shared subunits by the three sub-complexes, immunoprecipitated all the subunits found in the three sub-complexes (Figure 1B and Supplemental Figure S1E). Together, these immunoprecipitation data demonstrate the existence of three concurrently-expressed plant SWI/SNF family sub-complexes that have specific subunits and are separately assembled on a mutually exclusive catalytic subunit, BRM, SYD, or MINUs. We therefore termed these three plant SWI/SNF sub-complexes as BAS (<u>B</u>RM-<u>a</u>ssociated <u>S</u>WI/SNF complex), SAS (<u>S</u>YD-<u>a</u>ssociated <u>S</u>WI/SNF complex), and MAS (<u>M</u>INU-<u>a</u>ssociated <u>S</u>WI/SNF complex), respectively (Figure 1C).

### Comparison of SWI/SNF sub-complexes among yeast, human and Arabidopsis

SWI/SNF complexes are multi-protein machineries evolutionally conserved among yeasts, animals, and plants (Ho and Crabtree, 2010; Sarnowska et al., 2016). We compared the three distinct plant SWI/SNF complexes with the mammalian SWI/SNF complexes and identified several plant-specific properties in SWI/SNF complexes. First, in humans, the three different types of SWI/SNF sub-complexes, BAF, PBAF and ncBAF, shared the same ATPases (BRM/BRG1) and initial BAF core subunits (two SMARCC and one SMARCD). However, in *Arabidopsis*, SAS, MAS and BAS each used different ATPases and SMARCCs homologs. Specifically, the SAS complex contained SYD ATPase and SWI3D, the MAS complex included MINU1/2 ATPases and SWI3A/B, while BRM ATPase and SWI3C specifically presented in the BAS complex (Figure 2A). Second, plant SWI/SNF complexes uniquely lacked core, metazoan-conserved subunits such as SMARCE1 and pBRM1, but selectively evolved in plant-specific subunits that are not found in metazoan and yeast, including PSA1, PSA2, SHH2, SSIP1, SSIP2, and SSIP3 (Figure 2A). Third, the BAS complexes are equivalent to the mammalian ncBAF because they contain the identical paralog subunits (Yu et al., 2020; Yu et al., 2021). However, comparison of the subunit compositions of the SAS and MAS sub-complexes with those of human BAF complexes suggested that the SAS and MAS are plant-specific. Indeed, SAS complexes did not contain any PHD domain or bromodomain proteins that are signatures of mammalian PBAF and cBAF, but had three plant-specific proteins SSIP1/2/3 whose functions in the SAS remain to be investigated. In terms of MAS complexes, although they had PHD domain proteins TPF1/2 and OPF1/2 that are homologous to PHF10 subunits in PBAF and DPF1/2/3 in cBAF and contain BRD5 protein that is homologous to the PBAF-specific subunit BRD7, they lacked homologous subunits of PBRM1 and SS18, which are specialized subunits in PBAF and cBAF, respectively. In contrast, the MAS complexes contained several subunits (SHH2, PSA1, and PSA2) that are specifically presented in plants. Based on these comparisons, we propose that the SWI/SNF sub-complexes in different kingdoms came from an ancestral BAF complex. This ancestor BAF complex was firstly evolved into an ncBAF complex that has been preserved in three kingdoms by integrating the GLTSCR domain-containing subunits. Meanwhile, the ancestral BAF complex integrated different new subunits and discarded several original ones to evolve into the PBAF and cBAF sub-complexes in animals and fungi, as well as two plant-specific sub-complexes, SAS and MAS (Figure 2B).

**Figure 2.**
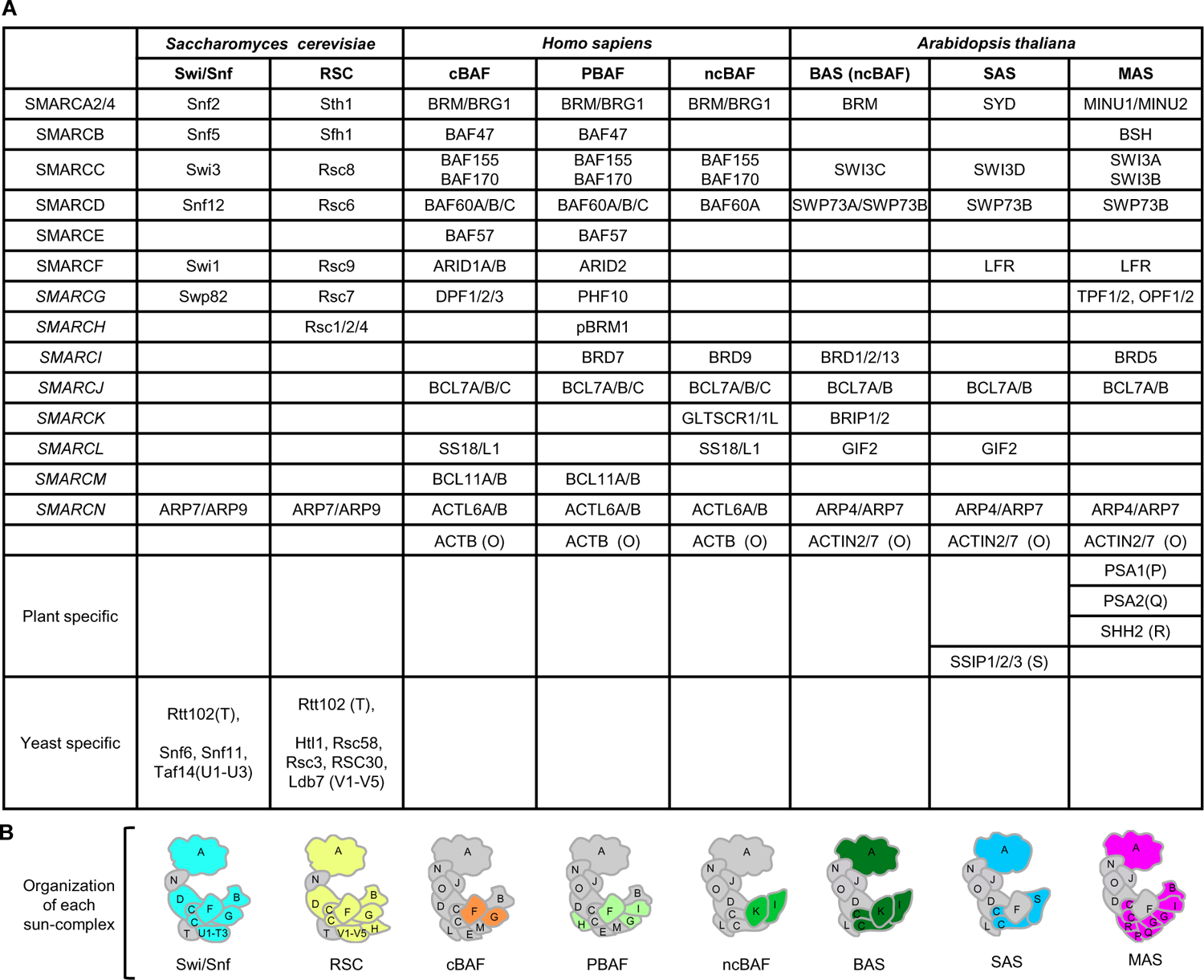
Comparison of subunit composition of SWI/SNF complexes in *Saccharomyces cerevisiae, Homo sapiens* and *Arabidopsis thaliana*. **A**, SWI/SNF sub-complex components in the three species. The *SMARC* names correspond to lineage-specific subunits previously identified (Hernández-García at al., 2022) and the characters in brackets represent different protein modules. **B**, Diagrams displaying the subunits composition of different SWI/SNF sub-complexes in eukaryotes. The specific subunits of each complex were highlighted by different colors, and the common subunits of different sub-complexes of the same species were marked with gray.

### Differential localization of the *Arabidopsis* SWI/SNF sub-complexes, SAS, MAS, and BAS on chromatin

To further characterize the three distinct plant SWI/SNF assemblies and to determine whether the differences of their subunit compositions may result in differential genomic targeting, we performed a comprehensive genome-wide mapping of SAS, MAS and BAS sub-complexes by performing chromatin immunoprecipitation-sequencing (ChIP-seq) using the stable expression transgenic plants, including pan-plant-SWI/SNF subunits (SWP73B and BCL7A/B) and complex specific subunits SYD and SWI3D for SAS, MINU2, SWI3A and BSH for MAS, and BRM, SWP73A and SWI3C for BAS (Supplemental Figure S2A). The ChIP-seq data of other BAS-specific subunits BRIP1/2 and BRD1/2/13 were previously described (Yu et al., 2020; Yu et al., 2021).

Consistent with biochemical results, subcomplex-specific subunits peaks comprised subsets of all pan-subunits peaks (Figure 3A and Supplemental Figure S2, A and B). The target genes of the three ATPases, SYD, MINU2 and BRM, showed significant overlaps (*n* = 4,093); however, they also exhibited specific binding sites, especially for MINU2 (*n* = 4,433) (Figure 3B). Similar results were shown for the other complex-specific subunits, such as SWI3D for SAS, SWI3A for MAS and SWI3C for BAS (Supplemental Figure S2C). Hierarchical clustering performed on ChIP-seq read density over the merged set of peaks across all ChIPs identified distinct, complex-specific enrichment on chromatin (Figure 3C and Supplemental Figure S2D). Indeed, SAS and MAS complexes were accumulated near transcription start sites (TSSs) relative to BAS complexes, which were substantially more enriched over gene bodies (Figure 3, D-E and Supplemental Figure S2E). Consistently, SYD and MINU2 located more frequently over the promoters of the target genes in comparison to BRM, which showed a preference for binding over exons and introns (Figure 3, F and G). Similarly, when we examined the enrichment patterns of other complex-specific subunits, we found that they exhibited similar distributions with their corresponding ATPases. For instance, the BAS specific subunits SWI3C, BRIP1/2 and SWP73A, like BRM, showed more enrichment over gene bodies compared with SWI3D and SWI3A (Supplemental Figure S2, F and G). Surprisingly, when comparing the binding summit of SWI3C with that of BRM, we found that the SWI3C summit was obviously shifted towards the TSSs, reflecting the different occupancies between SWI3C and the BRM ATPase within the BAS sub-complexes (Supplemental Figure S2, H and I). In this regard, one possible explanation is that the BAS sub-complexes contain three BRDs (BRD1/2/13) in their core module that can recognize histone acetylation on chromatin and a BRM in their ATPase module that may bind acetylated histones and DNA through its C-terminal bromodomain and AT-hook domain, respectively (Zhao et al., 2018; Yu et al., 2021). Supporting this notion, we found that the width of BAS complexes peaks was significantly larger than that of SAS and MAS complexes (Figure 3, E and H).

**Figure 3.**
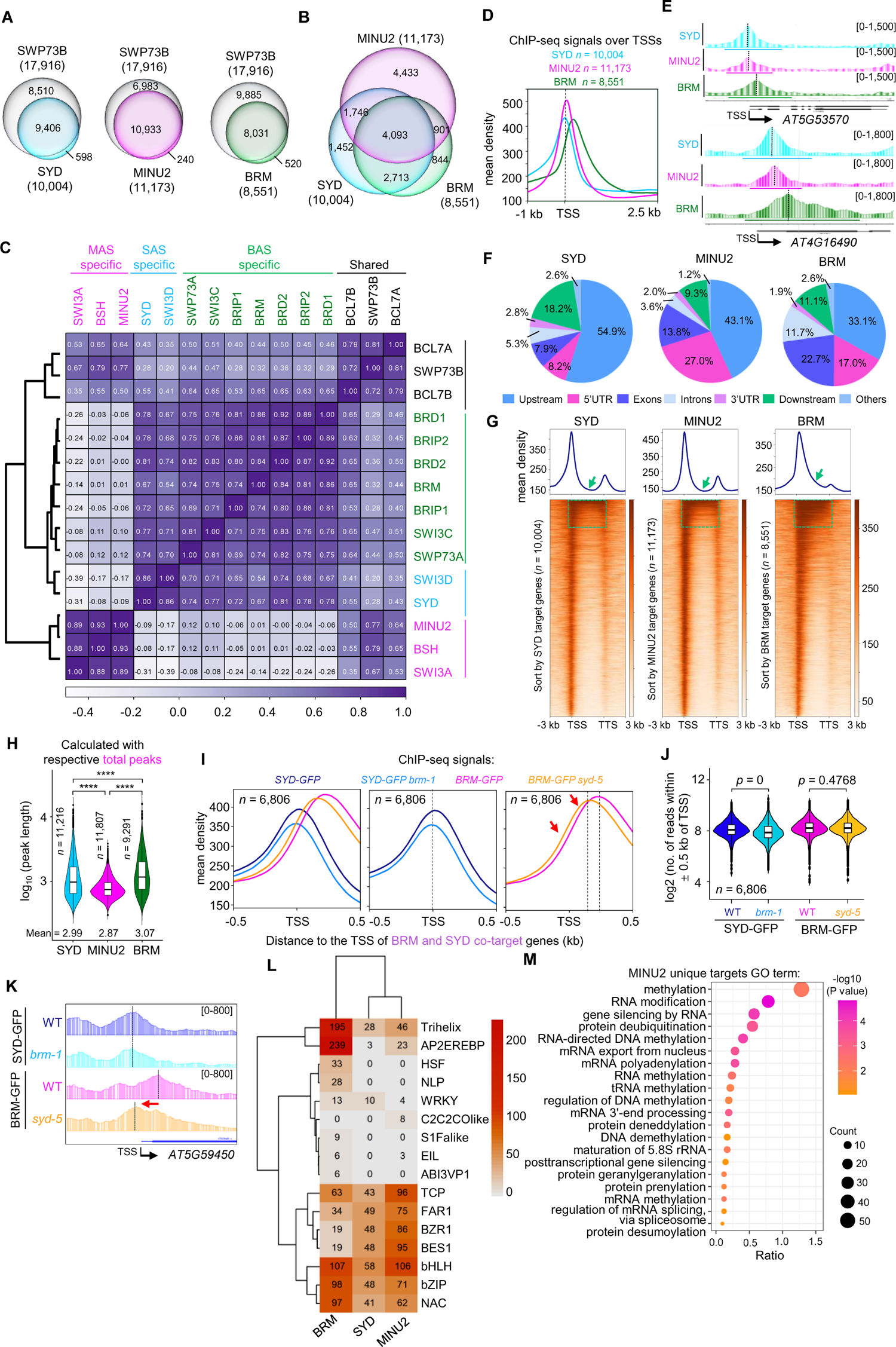
SAS, MAS and BAS sub-complexes show differential binding patterns in genome-wide. **A**, Venn diagrams displaying statistically significant overlaps among genes occupied by SYD, MINU2 or BRM with those by SWP73B. *P* values were calculated by the hypergeometric test. **B**, Venn diagram displaying statistically significant overlaps among genes occupied by SYD, MINU2 and BRM. *P* values were calculated by the hypergeometric test. **C**, Heatmap representing correlations between normalized ChIP-seq reads over a merged set of all SWI/SNF subunit peaks. **D**, SYD, MINU2, and BRM complex ChIP-seq read density distribution over the TSS and 2.5 kb into the gene body at their target genes. **E**, IGV views of ChIP-seq signals of SYD, MINU2 and BRM at representative genes. The dash lines indicted the peak summits. **F**, Pie charts showing the distribution of SYD, MINU2 and BRM peaks at genic and intergenic regions in the genome. **G**, Metagene plot and heatmap display the distribution of SYD, MINU2 and BRM peaks at genic and intergenic regions in the genome. Green arrows and dashed boxes indicate the gene body. **H**, Violin plots depicting the log_10_(peak length) of SYD, MINU2 or BRM at their corresponding peaks. *P* values are from two-tailed Mann-Whitney *U* test. **I**, Metagene plot showing the mean occupancy of SYD or BRM at their co-target genes. **J**, Violin plots depicting the log2 (no. of reads within ± 0.5 kb of TSS of BRM-GFP and SYD-GFP at their co-target genes, *P* values were determined by the two-tailed Mann-Whitney *U* test. **K**, IGV views of ChIP-seq signals of BRM and SYD at representative genes. The diagrams underneath indicate gene structure. The y-axis scales represent shifted merged MACS2 tag counts for every 10-bp window. **L**, Heatmap of CentriMo log adjusted p-values for top motifs returned by MEME-ChIP analysis for each ChIP-seq experiment. P-values were calculated using binomial test. The sequence covering 250 bp (SYD and MINU2) or 400 bp (BRM) on either side of each peak summit were used. **M**, Gene ontology analysis of MINU2 specific targets genes.

Next, we want to assess whether the three sub-complexes could mutually regulate their binding positions on chromatin. To this end, we examined the genomic occupancy of the BAS complexes upon the loss of the SAS, and vice versa. We introduced the loss-of-function *syd-5* mutant into the *BRM-GFP* transgenic line (*pBRM:BRM-GFP brm-1 syd-5*) and loss-of-function *brm-1* mutant into the *SYD-N-GFP* transgenic line (*pSYD:SYD-N-GFP syd-5 brm-1*) and then performed ChIP-seq assays. We found that the SYD occupancy density was decreased upon BRM mutation at the genes that are co-targeted by BRM and SYD, but the average occupancy intensity of BRM was almost the same in the *syd-5* background compared with WT (Figure 3, I-K). However, in the absence of SYD, the BRM binding position appeared to have a significant shift (from gene body to TSS) at BRM-SYD co-target genes (Figure 3, I and K), implying that SAS sub-complexes enable BAS sub-complexes to bind to chromatin accurately.

Motif analyses revealed a significant central enrichment of the three sub-complexes over known transcription factor motifs, including NAC, bZIP, bHLH, BES1, BZR1, FAR1 and TCP. However, BRM preferentially localized to Trihelix and AP2EREBP and specifically localized to HSF, NLP and ABI3VP1 (Figure 3L). In addition, MINU2 specifically localized to C2C2COlike (Figure 3L). These results imply specialized roles for SAS, MAS and BAS complexes at the corresponding motifs. Furthermore, Gene Ontology (GO) enrichment analysis of the three ATPases and core subunits (SWI3D, SWI3A and SWI3C) showed enrichment of shoot system development, response to light stimulus, growth, and flower development (Supplemental Figure S2J and Supplemental Figure S3). Of note, MAS sub-complex core members MINU2 and SWI3A were particularly enriched in many biological pathways different from BAS and SAS sub-complex, such as embryo development, mRNA processing, chromatin organization and DNA methylation (Figure 3M, Supplemental Figure S2J and Supplemental Figure S3). When we further analyzed the enrichment signals of SAS, MAS, and BAS on the genes involved in DNA methylation regulation, we found that MINU2 and SWI3A, the MAS-specific subunits, were significantly enriched on these genes, while the SAS and BAS showed less or no enrichment (Supplemental Figure S4). Collectively, these data suggested a potential specific role for the MAS sub-complexes in the regulation of DNA methylation.

We previously showed that BAS complexes subunits BRM and BRD1/2/13 preferentially occupied active histone modification markers (Zhao et al., 2018; Yu et al., 2021). In humans, all three BAF subcomplexes were localized to active enhancers and promoters (H3K27ac and H3K4me1) (Michel et al., 2018). When we analyzed the enrichment signals of the active markers (H3K9ac, H3K27ac, H4K5ac, H4K8ac, H3K4me2, H3K4me3, and H3K36me3) and the repressive markers (H3K27me3) at the peaks that are occupied by SYD, MINU2 or BRM, we found that the MINU2 and BRM binding peaks enriched by the active histone marks but are depleted of the repressive one (Supplemental Figure S5A). In contrast, the peaks center of SYD lacked the active markers but showed the highest H3K27me3 enrichment signals among the three ATPases (Supplemental Figure S5A). Moreover, SYD peaks displayed a weak Pol II enrichment relative to MINU2 and BRM peaks (Supplemental Figure S5A). When we repeated this analysis using the sub-complexes co-binding peaks and their unique binding peaks, we observed that the active markers were significantly enriched on the co-binding peaks of SYD-MINU2-BRM, as well as on the MINU2 unique and BRM unique peaks, but were almost not enriched on the SYD unique peaks (Supplemental Figure S5B and C). However, there is a strong enrichment of H3K27me3 signals on the SYD unique peak. Moreover, the same results were obtained when we used the complex specific subunit peaks for the analysis (Supplemental Figure S6). Together, these results demonstrate plant SWI/SNF complex-specific chromatin localization and indicate a specialized enrichment of SAS over the repressive chromatin regions, which has not been observed in human SWI/SNF complexes.

### SWI3D subunit acts in the SAS complexes to regulate genome-wide gene expression

Our biochemical data above suggested that among the four SMARCC paralogs in plants, SWI3D is selectively incorporated into the SAS complexes but not the MAS and BAS complexes (Figure 1). To determine whether SWI3D is functionally relevant to the SAS complexes, we compared the morphological phenotypes between *syd-5* and *swi3d-1* loss-of-function mutants and observed that the two mutant seedlings showed extraordinarily identical phenotypes, including root length, leaf shape, and plant growth status (Figure 4, A and B) (Sarnowski et al., 2005; Shu et al., 2021). Moreover, we introduced the *swi3d-1* into the *syd-5* and found that the *swi3d-1 syd-5* double mutants displayed the same phenotypes as *syd-5* and *swi3d-1* single mutants, showing that loss of SWI3D did not enhance the phenotypes of *syd-5* null mutant. Thus, SWI3D function selectively in the SAS complexes with SYD to regulate the plant development processes.

**Figure 4.**
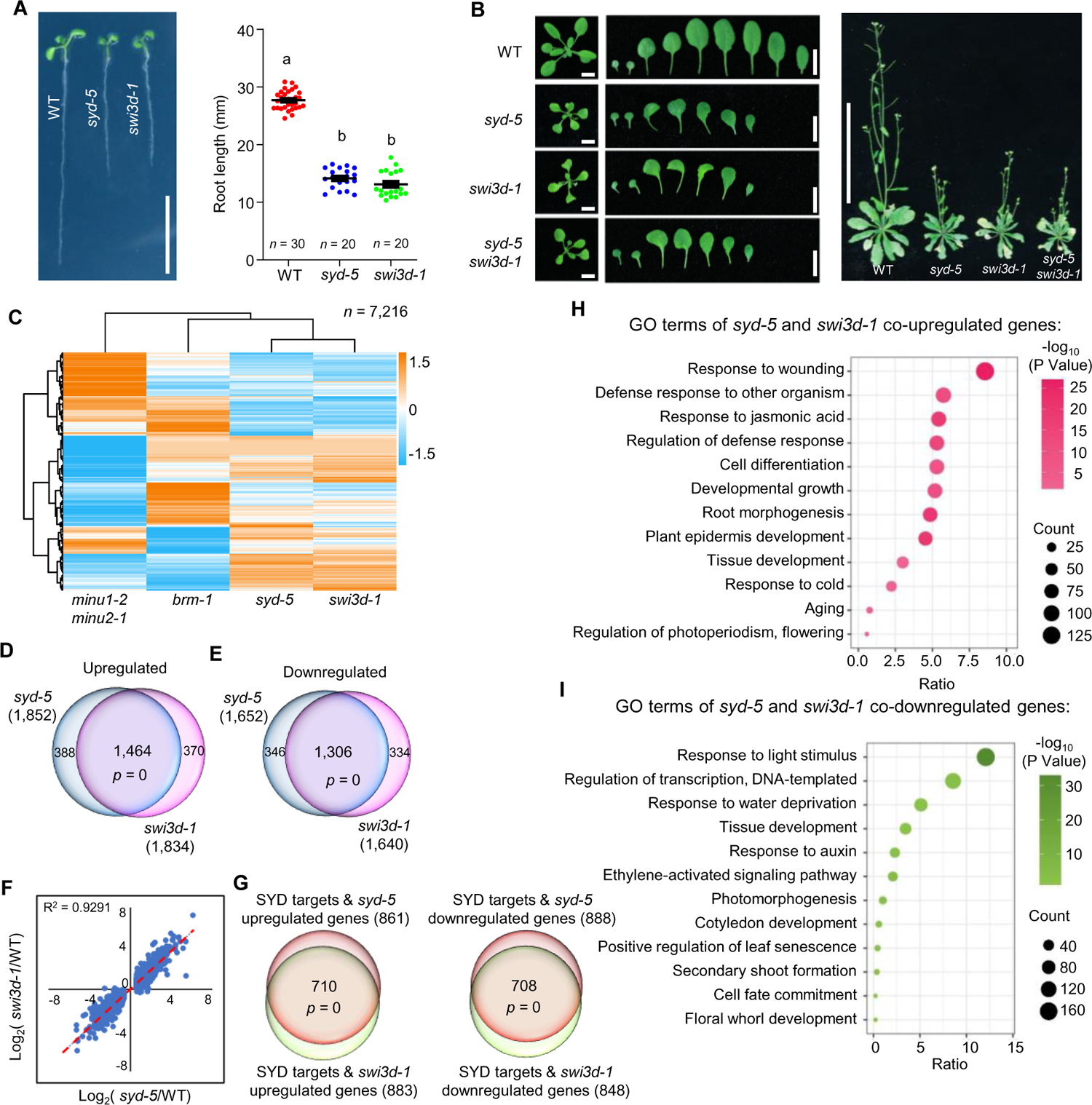
SWI3D and SYD co-regulate gene expression. **A**, root length of 7-day-old seedlings. Scale bars, 1 cm. Lowercase letters indicate significant differences between genetic backgrounds, as determined by the post hoc Tukey’s HSD test. The number of “n =” indicates the number of plants that were used. Data are presented as mean ± s.d.. **B**, Left, leaf phenotype of 21-day-old seedlings. Scale bars, 1 cm. Right, seedlings phenotype of 40-day-old seedlings. Scale bars, 10 cm. **C**, Heatmap showing hierarchical clustering of differentially expressed genes in different mutants (total mis-regulated genes, 7,216). **D-E**, Venn diagrams showing statistically significant overlaps between genes up- or downregulated in *syd-5* and those in *swi3d-1*. *P* = 0, hypergeometric test. **F**, Scatterplot of log2-fold change values over WT of *syd-5* versus *swi3d-1* at genes that were differentially expressed in *syd-5*. The line of best fit is shown in red, with adjusted *R* value indicated. Dots are mean values from three biologically independent experiments. **G**, Venn diagrams showing statistically significant overlaps between the mis-regulated genes targeted by SYD in *swi3d-1* and SYD directs regulated genes. *P* values were determined by hypergeometric test. **H-I**, Gene ontology analysis of genes co-up- or co-down-regulated in *syd-5* and *swi3d-1*.

To corroborate these findings, we analyzed RNA-sequencing (RNA-seq) data comparing the transcriptome of *swi3d-1* mutants with that of *syd-5*, *brm-1*, and *minu1-2 minu2-1* mutant seedlings. This global transcriptional profiling revealed similar effects on gene expression between *swi3d-1* and *syd-5* mutants, whereas loss of BRM or MINU1/2 (BAS- and MAS-ATPase, respectively) resulted in discordant transcriptional effects (Figure 4C). Indeed, approximately 80% of the 1,834 upregulated and 1,640 downregulated genes in *swi3d-1* (1,462 and 1,306, respectively) exhibited the same direction of mis-regulation in *syd-5* mutants (Figure 4, D and E). The transcriptome of *swi3d-1* was strongly positively correlated with that of mutants *syd-5* (correlation coefficient R^2^ = 0.9291) (Figure 4F). Furthermore, genes mis-regulated in *syd-5* or *swi3d-1* significantly overlapped with SYD target genes (Supplemental Figure S7). Moreover, the SYD target genes misregulated in *swi3d-1* showed a very high overlap with those misregulated in *syd-5* mutants (Figure 4G). GO term analysis showed that genes regulated by SYD and SWI3D are involved in regulating development of tissues and organs and responding to wounding and auxin (Figure 4, H and I). Together, these data demonstrate that SWI3D and SYD are in the same complex to regulate gene expression in *Arabidopsis*.

### SWI3D is required for the assembly and integrity of SAS sub-complexes

In mammals, the assembly of the SWI/SNF complexes is initiated by the formation of a “core module” that is constituted by two SMARCC and one SMARCD subunits. This initial trimer then acts as a platform for docking of other subunits for assembly toward fully formed SWI/SNF complexes (Mashtalir et al., 2018). Because Arabidopsis SWI3D is a paralog of SMARCCs in mammals (Figure 2A), we thought to biochemically evaluate the effects of SWI3D loss on SAS protein complex assembly and integrity. To this end, we introduced the *SYD-GFP* transgene into the *swi3d-1* mutant background and assessed the abundance and integrity of SAS complexes using immunoprecipitation. The loss of SWI3D did not significantly change the messenger RNA levels of *SYD-GFP* (Figure 5A) but resulted in substantially reduced protein abundance of SYD-GFP (Figure 5, B-D). Furthermore, immunoprecipitation of SYD-GFP followed by mass spectrometry analysis showed that loss of SWI3D resulted in significantly reduced peptides corresponding to SYD and near-complete degradation of SAS sub-complex components (Figure 5, E and F). Notably, the protein level of BRM, the BAS ATPase, showed no change in the *swi3d-1* background compared to WT (Supplemental Figure S8), confirming the specific downregulation of the SAS integrity by SWI3D mutation. Together, these data demonstrate a crucial role for the core module subunit SWI3D in the stabilization of the SAS sub-complexes and imply that SWI3D may be a part of the core module that is also necessary for the assembly of SWI/SNF complexes in *Arabidopsis*.

**Figure 5.**
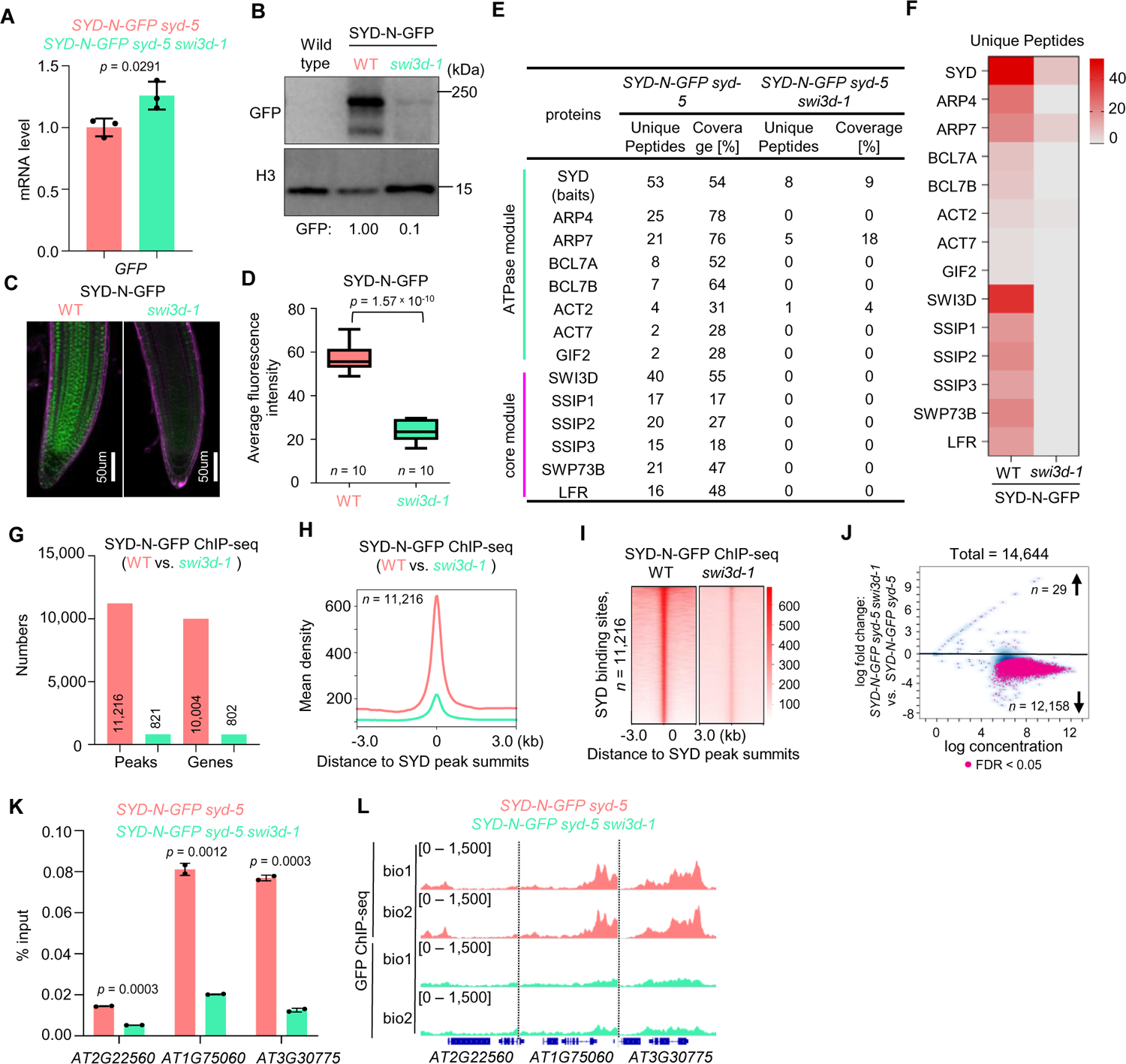
SWI3D is essential in maintaining SAS sub-complex integrity. **A**, The mRNA levels of *SYD-N-GFP* were determined by RT-qPCR in WT and *swi3d-1* background. *ACTIN2* was amplified as an internal control. Error bars are presented as mean values ± s.d. from three biological replicates. **B**, Immunoblot analysis showing the relative protein levels of SYD-N-GFP in WT and *swi3d-1*. The numbers at the bottom represent amounts normalized to the loading control, histone H3. WT was used as a GFP-free control. **C**, Confocal images of root tips showing nuclear localization of the SYD-N-GFP fusion protein in a WT and a *swi3d-1* background, respectively. The red fluorescent signal is derived from propidium iodide staining. **D**, Box plot showing the average fluorescence intensity of SYD-N-GFP in WT and *swi3d-1* mutants. The “*n =*” indicates the number of roots used. The boxes indicate the first and third quartiles, and the lines in the boxes indicate median values. Significant differences were determined by unpaired, two-tailed Student’s *t*-test. **E**, Unique peptide numbers of SAS sub-complex subunits identified by IP-MS in *SYD-N-GFP* under WT and *swi3d-1* background. **F**, Heatmap showing the unique peptides values of representative SAS sub-complex subunits identified by IP-MS in *SYD-N-GFP* under WT and *swi3d-1* background. **G**, Number of SYD binding sites (number of peaks or genes) in the WT and *swi3d-1* background. **H** and **I**, Metagene plot (**H**) and heatmap (**I**) represented the mean density of SYD occupancy at all SYD-occupied sites in *swi3d-1* compared with WT. The average SYD binding signals within 3 kb genomic regions flanking SYD peak summits were shown. **J**, Fold change (log2) in SYD occupancy between WT and *swi3d-1* background. Occupancy changes with false discovery rate (FDR) < 0.05 are highlighted. FDR values are multiple test-corrected Wilcoxon test *P* values, two biological replicates. **K**, ChIP-qPCR validation of SYD occupancy at representative target genes using ChIP DNA samples independent from those used for ChIP-seq. The *AT2G22560* was used as the negative control. Error bars are presented as mean values ± s.d. from two biological replicates. Unpaired, two-tailed Student’s *t*-test. **L**, IGV views of SYD occupancy at selected loci in the WT and *swi3d-1* background. The *y*-axis scales represent shifted merged MACS2 tag counts for every 10 bp window.

We next performed ChIP-seq assays comparing the genome-wide occupancy of SYD in WT with that in *swi3d-1* to assess the impact of SWI3D loss on the genome-wide targeting of the SAS complexes. In line with our biochemical findings, we observed substantial attenuation in SAS complex occupancy on the genome. The numbers of SYD-associated genomic sites and corresponding genes were extremely decreased in *swi3d-1* compared to WT (Figure 5G). Consistently, the occupancy of SYD on the target genes was nearly disrupted in *swi3d-1* (Figure 5, H and I). Furthermore, at more than 90% of SYD binding sites, we found a marked reduction in, or elimination of, SYD occupancy in the absence of SWI3D, while only 29 loci showed an increase (Figure 5J). Independent ChIP-qPCR confirmed the reduction of SYD occupancy at individual loci in *swi3d-1* (Figure 5, K and L). These data indicate that SWI3D loss disrupts SYD occupancy on chromatin and target gene expression.

### The SANT and SWIRM assoc_1 domain of SWI3D are required for the stability of SAS sub-complexes

We next sought to identify specific regions on SWI3D that uniquely underlie its function in SAS complex stability maintenance. SWI3D contains five conserved domains, including SWIRM, ZnF, SANT, RPT1 and SWIRM_assoc_1. We generated individual domain-truncated SWI3D fragments tagged by GFP and stably expressed them in the *swi3d-1* mutant background (Figure 6A). Deletion of the SWIRM, ZnF and PRT1 domains did not affect the ability of SWI3D to rescue the *swi3d-1* mutant phenotypes (Figure 6B), indicating that they are dispensable for SWI3D, at least under our growth conditions. In contrast, the deletion of SWIRM_assoc_1 caused a complete failure to restore the morphological defects of *swi3d-*1, while SWI3D truncation lacking the SANT domain only partially recovered the phenotypes (Figure 6B). These data highlight the importance of the SANT and SWIRM_assoc_1 domains of SWI3D in SAS complexes function.

**Figure 6.**
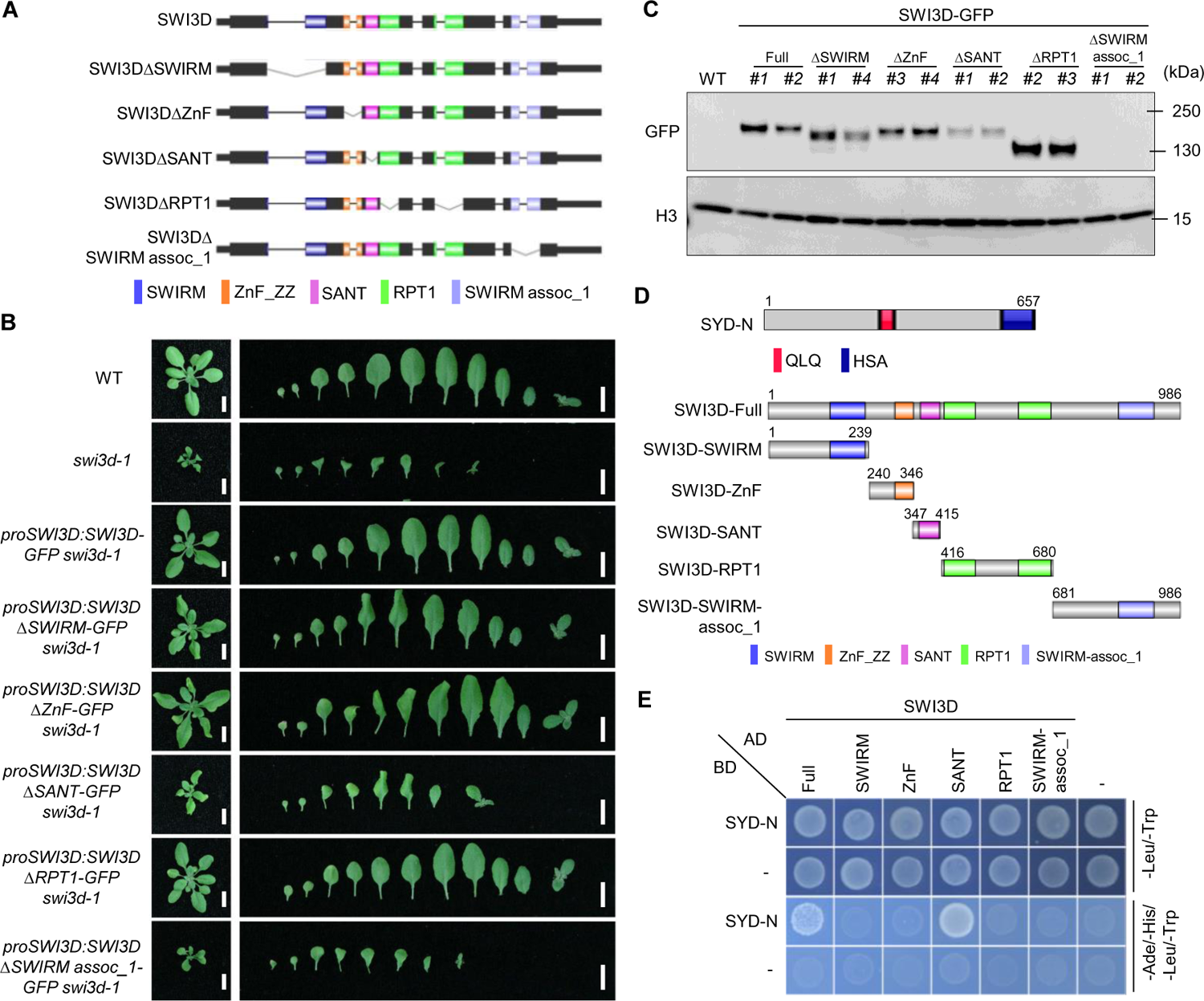
The SANT and SWIRM assoc-1 domain are required for maintaining the stability of SWI3D protein. **A**, Schematic illustration of the SWI3D protein and its truncated versions. **B**, Leaf phenotype of 21-day-old seedlings. Scale bars, 1 cm. **C**, Western blot analysis using an anti-GFP antibody shows the accumulation of SWI3D protein and its truncated versions. For each plot, the antibody used is indicated on the left, and the sizes of the protein markers are indicated on the right. H3 serves as a loading control. **D**, Schematic illustration of the the N-terminal of SYD protein, SWI3D and the different truncated versions of SWI3D. **E**, Y2H assay to examine the direct interaction between SYD and SWI3D. Growth of transformed yeast on permissive SD^-Ade-His-Leu-Trp^ medium indicate interaction.

To understand the role of SANT and SWIRM_assoc_1 in the SAS complexes, we examined the protein levels of the truncated SWI3Ds. We observed that the deletion of the SANT domain resulted in a reduction in the level of SWI3D protein, and the deletion of the SWIRM_assoc_1 domain led to a failure to accumulate SWI3D protein (Figure 6C). Notably, the decrease in their protein contents was not caused by the reduction of their transcription levels (Supplemental Figure S9A). Interestingly, the SANT and SWIRM assoc_1 domains are evolutionary conserved in different eukaryotes, suggesting that their role in modulating SWI3D protein abundance could possibly be a conserved function (Supplemental Figure S9B). To further elaborate on this, we performed yeast two-hybrid (Y2H) assays to test which domain(s) of SWI3D interacts with the SYD (Figure 6D). We found that only the SANT domain, but not the other domains, is required for interaction with SYD ATPase and serves as a SAS binding domain (Figure 6E). Together, these results demonstrate that the SWI3D SANT domain is a SAS-complex-binding domain that underlies the critical role of SWI3D in the assembly of the SAS complexes.

### The SYD ATPase module is required for the stability of the core module of the SAS sub-complex

Our data so far showed that the core module subunit SWI3D of SAS complexes plays a vital role in regulating the integrity of the complex. In mammals, recent studies indicated that loss of ATPases results in a specific disruption of the ATPase module but has no effect on the assembly of the core module (Mashtalir et al., 2018; Pan et al., 2019). However, our above data showed that in Arabidopsis, disruption of SWI3D-SYD interaction by deleting the SWI3D SANT domain leads to a decrease in SWI3D protein abundance. We therefore speculated that, in contrast to their human paralogs, ATPase subunits in plants might be required to stabilize the core module.

To test this possibility, we introduced the loss-of-function *syd-5* mutant into the *SWI3D-GFP* transgenic line. Surprisingly, we found that the SWI3D-GFP protein levels were dramatically decreased in the absence of the SYD ATPase (Figure 7A); however, the expression level of *SWI3D-GFP* mRNA did not change significantly (Figure 7B). Furthermore, we conducted IP-MS experiments and found that in *syd-5* mutants, both the core module subunits (such as SWP73B and LFR) and the ATPase module subunits (such as ARP4/7 and BCL7A/B) were not co-immunoprecipitated (Figure 7C). These results indicate that the ATPase in SAS sub-complexes play an essential role in the stability of the core module. To test whether the requirement of ATPase for the stability of the complex is not restricted to SAS complexes but a widespread phenomenon for plant SWI/SNF complexes, we examined the consequences of BRM loss on the stability of SWI3C core module subunit in BAS complexes. We introduced the loss-of-function *brm-1* mutant into the *SWI3C-GFP* transgenic line and performed IP-MS assays. Upon the loss of BRM, SWI3C barely immunoprecipitated ATPase module subunits ARP4/7 and BCL7A/B and less effectively immunoprecipitated the core subunit BRIP1/2, BRD1/13 and SWP73B (Supplemental Figure S10). Together, these data demonstrate that the ATPase module enables the full assembly of the core module of Arabidopsis SWI/SNF complexes, a plant-specific feature distinct from animal SWI/SNFs.

**Figure 7.**
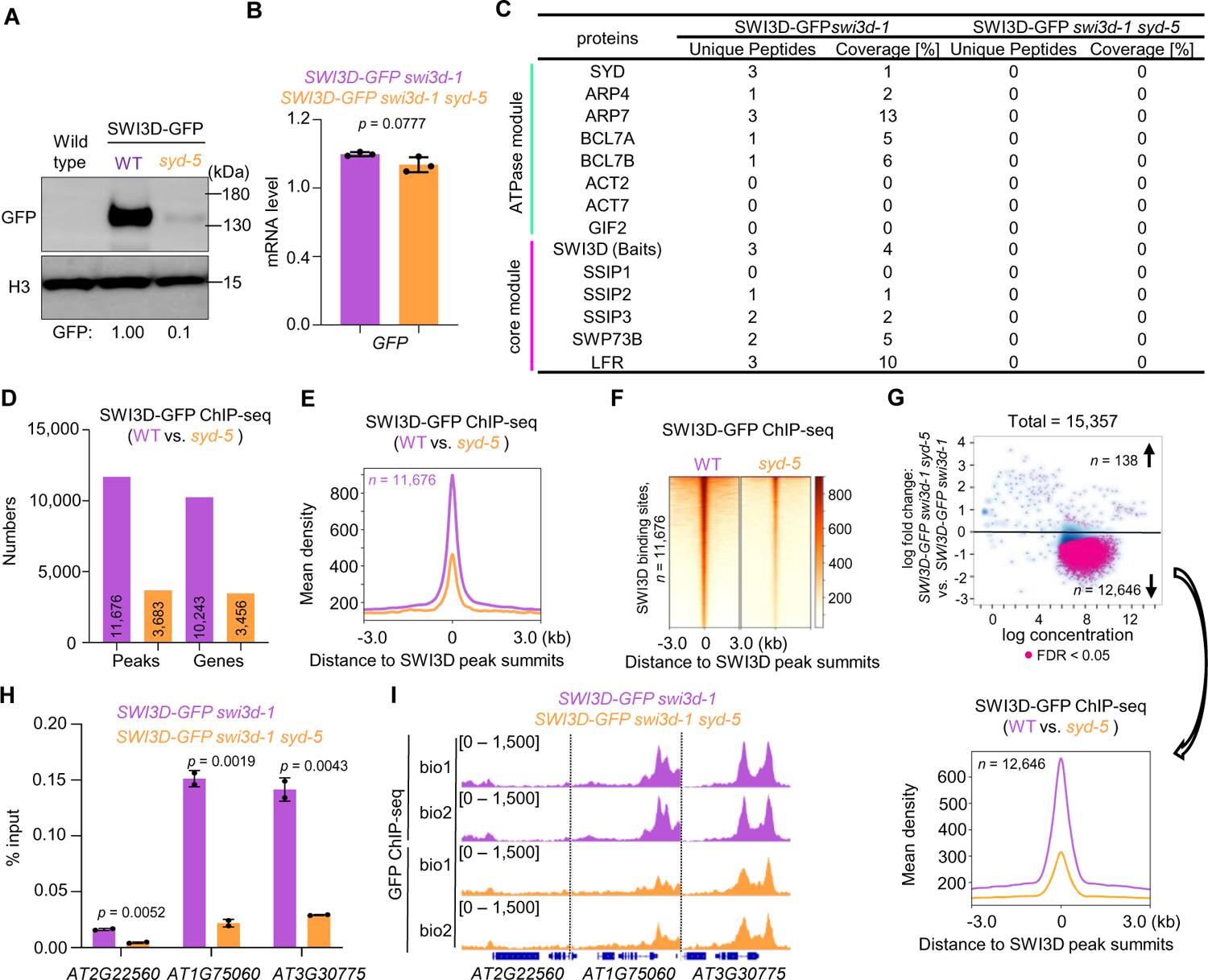
The ATPase module is required for stability of the core module. **A**, Immunoblot analysis showing the relative protein levels of SWI3D-GFP in WT and *syd-5* background. The numbers at the bottom represent amounts normalized to the loading control, histone H3. WT was used as a GFP-free control. **B**, The mRNA levels of SWI3D-*GFP* were determined by RT-qPCR in WT and *syd-5* background. *ACTIN2* was amplified as an internal control. Error bars are presented as mean values ± s.d. from three biological replicates. **C**, Unique peptide numbers of SAS sub-complex subunits identified by IP-MS in *SWI3D-GFP* under WT and *syd-5* background. **D**, Number of SWI3D binding sites (number of peaks or genes) in the WT and *syd-5* background. **E** and **F**, Metagene plot (**E**) and heatmap (**F**) represented the mean density of SWI3D occupancy at all SWI3D-occupied sites in *syd-5* compared with WT. The average SWI3D binding signals within 3 kb genomic regions flanking SWI3D peak summits were shown. **G**, Fold change (log2) in SWI3D occupancy between WT and *syd-5* background. Occupancy changes with false discovery rate (FDR) < 0.05 are highlighted. FDR values are multiple test-corrected Wilcoxon test *P* values, two biological replicates per ChIP. **H**, ChIP-qPCR validation of SWI3D occupancy at representative target genes using ChIP DNA samples independent from those used for ChIP-seq. The *AT2G22560* was used as the negative control. Error bars are presented as mean values ± s.d. from two biological replicates. Unpaired, two-tailed Student’s *t*-test. **I**, IGV views of SWI3D occupancy at selected loci in the WT and *syd-5* background. The *y*-axis scales represent shifted merged MACS2 tag counts for every 10 bp window.

Finally, we tested the genomic binding of SWI3D in the *syd-5* mutants. ChIP-seq results showed that compared with the enrichment signals of SWI3D in the wild-type background, the number and the signal intensity of SWI3D binding peaks in the genome decreased sharply in *syd-5* (Figure 7, D-G). Independent ChIP-qPCR confirmed the reduction of SWI3D occupancy at individual loci in *syd-5* mutant (Figure 7, H and I). However, a subset of genomic sites was still occupied by SWI3D in the absence of SYD ATPase (Figure 7D, E). Interestingly, we found that the SYD-independent SWI3D binding sites were more accessible than the SYD-dependent SWI3D binding sites (Supplemental Figure S11), possibly underscoring the different dependencies on the SYD ATPase in SAS genomic targeting.

## Discussion

In this study, we demonstrate that Arabidopsis has three distinct SWI/SNF sub-complexes (SAS, MAS, and BAS), each containing complex-specific subunits that probably define their identity (Figures 1 and 2). We also comprehensively compared the chromatin binding profiles for the three sub-complexes relative to defined chromatin features (Figure 3). The SAS, MAS, and BAS sub-complexes showed differential occupancy patterns on target genes, associated unique cis-motifs, and localized to chromatin with different histone modifications, possibly underlying complex-specific functions. Moreover, the loss of SAS complexes results in a unexcepted shift of the BAS occupancy from gene body to TSSs, providing new insights into our understanding of how the genomic targeting of different SWI/SNF sub-complexes is coordinated for chromatin structure regulation. Finally, our work found a mutual dependency of the core module and the ATPase module of SAS in their assembly for maintaining the integrity of the complex in plants.

In the past two decades, numerous investigations on plant SWI/SNF complexes subunits have revealed various phenotypes caused by the perturbation of the subunits. However, due to the lack of information regarding to the subunit composition of plant SWI/SNF complexes, our understanding to the mechanisms by which these subunits regulate plant development and responses to environment stimulus has been severely affected. Moreover, in vitro protein-protein interaction assays may not identify endogenous, physiologically relevant complex-specific subunits. For example, previous yeast-two-hybrid assays showed the interaction between SWI3D and BRM (Bezhani et al., 2007). Here, our biochemical data demonstrated that SWI3C, but not SWI3D, is a BAS-specific subunit (Figure 1), which elaborates the similar phenotypes between *swi3c* and *brm* mutants, rather than between *swi3c* and *syd* or *minu* mutants (Wagner D, 2002; Archacki et al., 2009; Sang et al., 2012). Consistently, we found that the phenotype and transcriptome of *swi3d-1* are reminiscent of *syd-5*, in support of the notion that SWI3D and SYD are in the SAS complexes. Moreover, the *swp73a* single mutant showed no phenotypes compared with the wild-type, but the *swp73b* single mutation caused severe alterations in the development of vegetative and reproductive organs (Sacharowski et al., 2015; Huang et al., 2021). Our data showed that SWP73B exists in all three SWI/SNF sub-complex, but SWP73A only belongs to the BAS sub-complexes. Thus, in the *swp73a* mutant, SWP73B could act in the three sub-complexes to maintain plant growth and development. However, in the absence of SWP73B, both SAS and MAS sub-complexes were disintegrated, explaining the strong developmental disorders. Overall, our data underscore the importance of comprehensively defining the subunits composition of the complexes to illustrate subunit functions within the context of sub-complexes.

One particularly unexpected result is that the ATPase module (SYD or BRM) is required for global complex stability and the interaction of core module subunits in SAS and BAS complexes in Arabidopsis. Previous biochemical and structural studies in humans showed that the mammalian ATPase module is the last to be incorporated into SWI/SNF complexes and therefore not required for the core module assembly (Mashtalir et al., 2018; Michel et al., 2018; Pan et al., 2019). Thus, our studies suggest an alternative, plant-specific assembly pathway in SWI/SNF complexes, further suggesting the separation and divergence of SWI/SNF complexes in eukaryotes.

Intriguingly, PAS1, PAS2, SHH2 and SSIP1/2/3 exist as conversed, plant-specific SWI/SNF subunits, suggesting an evolutionarily conserved function for those subunits that could not be appreciated by conventional sequence conservation analyses. Future functional and mechanistic characterization of those subunits would provide novel insight into the roles of plant SWI/SNF complexes in chromatin regulation. Furthermore, compared to SAS and BAS, the MAS sub-complexes showed a preferential occupancy at genes involved in the regulation of DNA methylation (Figure 3M, Supplemental Figure 3 and 4). Previous studies have indicated that SWI3B, which we found to be a MAS-specific subunit, acts to promote DNA methylation at a subset of genomic regions (Yang et al., 2020). Moreover, the MAS complexes contain an SHH2 subunit, whose paralog, SHH1, functions as a core component of RNA-directed DNA methylation by assisting in the recruitment of Pol IV (Law et al., 2013; Zhang et al., 2013). Together, these observations imply that the MAS sub-complexes may play a specialized role in regulating DNA methylation, which warrants future investigation.

A recent study demonstrated that the abundance of SWI/SNF complexes in human cells is not static but can be dynamically altered as a response to environmental changes (Tran et al., 2022). For example, the protein levels of PBAF-specific subunits ARID2 and PBRM1, and of ncBAF-specific subunit BRD9 are downregulated in response to hypoxic stresses, while all cBAF-specific members are retained. Here, our GO analysis of the three plant SWI/SNF sub-complexes revealed varying degrees of enrichments in defense response to bacterium, response to salt stress, immune system process, response to light stimulus and cellular response to phosphate starvation and so on. Thus, we speculate that the abundances of the three plant SWI/SNF sub-complexes may be dynamic rather than static in response to multifarious environmental signals for timely and precisely regulation of gene expression.

Finally, our proteomic analysis showed that some homologous subunits in SAS, MAS, or BAS are mutually exclusive in the complexes. For example, for SAS complexes, the three SSIPs (SSIP1, SSIP2, SSIP3) are separately incorporated into the SAS complexes (Figure 1B). Similarly, OPF1, the MAS-specific subunit, did not exist in the OPF2-containing MAS complexes (Diego-Martin et al., 2022). Finally, BAS-specific subunit BRIP1 cannot co-immunoprecipitated with BRIP2, and BRD1 cannot co-immunoprecipitated with BRD2 and BRD13 (Figure 1B). These observations suggest that the existence of multiple subunit paralogs across these three distinct SWI/ SNF sub-complexes may result in further diversification. Based on the multiple subunit paralogs across the three SWI/SNF sub-complexes, we calculated the full set of possible combinations (Supplemental Figure S12), in which there are at least 36 possible combinations of SAS sub-complexes, 288 possible combinations of MAS sub-complexes, and 144 possible combinations of BAS sub-complexes.

In summary, the illustration of three plant SWI/SNF complexes by our work provides a critical foundation for further structural and functional characterization of this family of plant chromatin remodeling complexes that play crucial roles in regulating plant development and signaling responses through chromatin modulation. Our results highlight and reinforce the power of examining the organization, assembly and genomic targeting to advance our mechanistic understanding of SWI/SNF-mediated chromatin remodeling in plants.

## MATERIALS AND METHODS

### Plant materials and growth conditions

Transfer DNA insertion lines, *minu2-1* (SALK_057856), *swi3a-3* (SALK_068234), *bsh* (SALK_058513), *syd-5* (SALK_023209), *swi3d-1* (SALK_100310), *swp73a-2* (SALK_083920), *swp73b-1* (SALK_113834), *ssip2-1* (SALK_109947), *ssip3-1* (SALK_010574), *bcl7a-1* (SALK_027934) and *bcl7b-1* (SALK_029285) were obtained from the Arabidopsis Biological Resource Center (ABRC). Mutants *brm-1* (SALK_030046), *pBRM:BRM-GFP brm-1* and *pSYD:SYD-N-GFP syd-5* transgenic plants were previously described (Li et al., 2016; Shu et al., 2021). Primers used for genotyping are listed in Supplementary Table 1.

For RT-qPCR/RNA-seq, ChIP-qPCR/ChIP-seq and IP-MS assays, *Arabidopsis* seeds were sterilized with 20% sodium hypochlorite solution for 15 mins, washed with sterile water five times, and then stratified at 4 °C in darkness for three days. Seeds were then sown on ½ Murashige and Skoog (MS) medium containing 1% sucrose and 0.6% agar. For phenotypic analysis, seeds were sown on a mixture of soil and vermiculite (1:1). Seedlings were grown under long-day conditions (16 h light/8 h dark) at 22 °C.

### Generation of transgenic plants

Full-length genomic regions of *SWI3A*, *SWI3B*, *SWI3C*, *SWI3D*, *SWP73A*, *BSH*, *MINU2*, *SSIP1*, *SSIP2*, *SSIP3*, *SWI3DΔSWIRM*, *SWI3DΔZnF*, *SWI3DΔSANT*, *SWI3DΔRPT1* and *SWI3DΔSWIRM_assoc_1,* driven by their native promoters were cloned into *pCAMBIA1302* vector by using ClonExpress Ultra One Step Cloning Kit (Vazyme, Cat. No. C115-01) and ClonExpress MultiS One Step Cloning Kit (Vazyme, Cat. No. C113-01). Full-length genomic regions of *BCL7A*, *BCL7B* and *SWP73B*, driven by their native promoters were amplified from genomic DNA by PCR, then cloned into the *pDONR221* vector by BP reaction (Invitrogen), and further subcloned into the destination plasmid *pMDC107* (Curtis and Grossniklaus, 2003) by LR reaction (Invitrogen). The constructs were introduced into *Agrobacterium tumefaciens* strain *GV3101* and were then used to transform corresponding single mutant or WT (e.g., SWI3B, SWI3C and SSIP1) plants using the floral dip method (Clough and Bent, 1998). To obtain *BRM-GFP brm-1 syd-5* and *BRM-GFP brm-1 swi3d-1* transgenic plant, the *BRM-GFP brm-1* transgenic plant was crossed with *syd-5* and *swi3d-1* single mutant. Similar like this, the *SYD-N-GFP syd-5* transgenic plant was crossed with *brm-1* single mutant to obtain *SYD-N-GFP syd-5 brm-1* transgenic plant, the *SYD-N-GFP syd-5* transgenic plant was crossed with *swi3d-1* single mutant to obtain *SYD-N-GFP syd-5 swi3d-1* transgenic plant, the *SWI3C-GFP* transgenic plant was crossed with *brm-1* single mutant to obtain *SWI3C-GFP brm-1* transgenic plant and the *SWI3D-GFP swi3d-1* transgenic plant was crossed with *syd-5* single mutant to obtain *SWI3D-GFP swi3d-1 syd-5* transgenic plant. Primers used for constructing are listed in Supplementary Table 1.

### Y2H assay

The full-length or truncated coding regions of SYD or SWI3D were cloned into *pGADT7* (AD) or *pGBKT7* (BD). Then the AD and BD plasmids were co-transformed into the yeast strain *AH109* and spread on the medium that lacking leucine (Leu) and tryptophan (Trp) (SD-Leu/-Trp). Positive colonies on SD-Leu/-Trp medium were further picked up and dropped on the selection medium lacking adenine (Ade), histidine (His), Leu and Trp (SD-Leu/-Trp/Ade/His). Primers used for constructing are listed in Supplementary Table 1.

### Confocal microscopy

The GFP signals from the root tips of transgenic plants were observed using the LSM880 microscope. The average fluorescence intensity of the GFP signals were calculated by the Histo function of the LSM880 microscope.

### Co-immunoprecipitation and mass spectrometry (IP-MS)

For IP-MS, about 5 g of 14-day-old seedlings under long-day conditions were harvested and ground to a fine powder in liquid nitrogen. Then, the powers were collected and homogenized in 10 ml of lysis buffer (50 mM HEPES [pH 7.5], 300 mM NaCl, 10 mM EDTA, 1% Triton X-100, 0.2% NP-40, 10% glycerol, 2 mM DTT, 1× Complete Protease Inhibitor Cocktail (Roche)) at 4℃ for 30 min. After centrifugation at 11,000 rpm and 4°C for 15 min (twice), the supernatant was diluted by equal volume dilution buffer (50 mM HEPES [pH 7.5], 10 mM EDTA, 10% glycerol, 2 mM DTT, 1× Complete Protease Inhibitor Cocktail (Roche)) and then incubated with GFP_trap beads (Cat. No. KTSM1301) at 4°C for 3 h with gently rotation. Beads were then washed three times with washing buffer (50 mM HEPES [pH 7.5], 150 mM NaCl, 10 mM EDTA, 0.2% Triton X-100, 0.1% NP-40, 10% glycerol). Proteins were eluted in SDS loading buffer and incubated at 55℃ for 10 min, followed by immunoblotting or silver staining.

For mass spectrometry, the immunoprecipitated proteins were eluted using 0.2 M glycine solution (pH 2.5), and then subjected to reduction with dithiothreitol, alkylation with iodoacetamide and digested with trypsin (Thermo Fisher, Cat. No. 90057, MS grade). The samples were analyzed on a Thermo Scientific Q Exactive HF mass spectrometer. Spectral data were searched against the TAIR10 database using Protein Prospector 4.0. Two or three biological replicates were included in the IP-MS analysis. Raw data were searched against the TAIR10. Default settings for Label-free quantitation (LFQ) analysis using MaxQuant (Tyanova et al., 2016a) and Perseus (Tyanova et al., 2016b) software were applied to calculate the LFQ intensities with default settings.

### Nuclear protein extraction

For nuclear protein extraction, 0.2 g of 14-day-old seedlings grown on 1/2 MS medium was ground to fine powder in liquid nitrogen to extract the nucleoproteins. Nuclei were isolated following the ChIP protocol (Li et al., 2016) without tissue fixation. Briefly, nuclear proteins were released by incubating the nuclei preparation in 200 ml of lysis buffer (50 mM Tris-HCl, 10 mM EDTA, 1% SDS, and 1× protease inhibitors) for 3h at 4℃. Then, the extract was diluted with equal volume of ChIP dilution buffer (16.7 mM Tris-HCl [pH 8.0], 167 mM NaCl, and 1.1% Triton X-100) and centrifuged at 15,000 g for 10 min at 4℃ to remove debris, followed by immunoblotting.

### Immunoblotting

Proteins were loaded onto 4%-20% gradient protein gels (GenScript, SurePAGE, Cat. No. M00655) and 4%-20% Precast Protein Plus Gel (Yeasen, Cat. No. 36256ES10) at 120 V for 2 h. A wet transformation was performed at 90 V for 90 min in ice-cold transfer buffer. After that, the membranes were blocked in 5% non-fat milk at room temperature for 1 h on a shaking table (60 rpm). Finally, the blocked membranes were incubated in the corresponding antibodies solutions at room temperature for another 3h. The following antibodies were used: anti-GFP (Abcam, Cat. No. ab290, 1:10,000 dilution), anti-H3 (Proteintech, Cat. No. 17168-1-AP, 1:10,000 dilution). The intensities of blotting signals were quantified using ImageJ software (v.1.50i). Uncropped scans of immunoblotting results are shown in Supplementary Figure 13.

### Silver staining

For silver staining, samples were run on a 4%-20% gradient protein gels (GenScript, SurePAGE, Cat. No. M00655) and stained with Fast Silver Stain Kit (Beyotime, Cat. No. P0017S) according to the manufacturer’s instructions.

### RNA isolation, RT-qPCR and RNA-seq analyses

Total RNA was extracted from 14-day-old Arabidopsis seedlings grown under long-day conditions using the HiPure Unviersal RNA Mini Kit (MAGEN, Cat. No. R4130) according to the manufacturer’s instructions. FOR RT-qPCR, 1 μg RNA was used for DNase digestion and reverse transcription using HiScript III 1st Strand cDNA Synthesis Kit (+gDNA wiper) (Vazyme, Cat. No. R312-01). The transcribed cDNA underwent qPCR assays were performed using ChamQ Universal SYBR qPCR Master Mix (Vazyme, Cat. No. Q711-02) in the StepOne Plus (Applied Biosystems). Results were repeated with three biological replicates. Quantification was analyzed with the relative -ΔΔCt method (Livak and Schmittgen, 2001). *ACTIN2* were served as the control for mRNA analyses. The sequences of primers used are listed in Supplementary Table 1.

For RNA-seq analyses, RNA from three biological replicates were sequenced separately at Novogene (sequencing method: nova-seq PE150). After removal of adapters and low-quality reads, the clean reads were mapped to the TAIR10 Arabidopsis genome using TopHat (v2.1.1) with default settings (Kim et al., 2013), except that a minimum intron length of 20 base pairs (Snelders et al.) and a maximum intron length of 4,000 bp were required. Mapped reads were then assembled according to the TAIR10 version of genome annotation using Cufflinks (v.2.1.1) with default settings (Trapnell et al., 2012). For analysis of differential expression, the assembled transcripts from three independent biological replicates in WT and other mutants (*syd-5* and *swi3d-1*) were included and compared using Cuffdiff (v.2.1.1) with default settings (Trapnell et al., 2012). Finally, genes with at least 1.5-fold change in expression (P < 0.05) were considered differentially expressed (see Supplementary Table 2 for details). To calculate the significance of the overlap of two groups of genes drawn from the set of genes, the total number of genes in the Arabidopsis genome used was 34,218 (27,655 coding and 6,563 non-coding genes) according to EnsemblPlants (http://plants.ensembl.org/index.html).

### ChIP and ChIP-seq analysis

ChIP experiments were performed as previously described (Gendrel et al., 2005; Li et al., 2016; Yu et al., 2021) with minor changes. In brief, 14-day-old seedlings (0.5 g for each biological replicate) grown on ½-strength MS medium under long-day conditions were fixed using 1% formaldehyde under a vacuum for 15 min and then ground into a fine powder in liquid nitrogen. The chromatin was sonicated into 300-500 bp fragments using a Bioruptor sonicator with a 30/45-s on/off cycle (27 total on cycles). Immunoprecipitation was performed using 1 μl of anti-GFP (Abcam, Cat. No. ab290) plus 40 μl Agarose beads (Abcam, Cat. No. 16-157) at 4°C overnight. Next day, the beads were washed subsequently with low-salt buffer, high-salt buffer, ChIP wash buffer and TE buffer, and then the immunoprecipitated chromatin were eluted with Elution buffer. Finally, after the eluate subsequently subjected to reverse crosslinks, RNase digestion and Proteinase K digestion, the DNA was purified by phenol/chloroform/isoamyl. ChIP-qPCR was performed with three biological replicates, and the results were calculated as a percentage of input DNA according to the Champion ChIP-qPCR user manual (SABioscience). The primers used for ChIP-qPCR are listed in Supplementary Table 1.

For ChIP-seq, 1 g of seedlings was used, and the ChIPed DNA was purified using the MinElute PCR purification kit (Qiagen, Cat. No. 28004). Libraries were constructed with 2ng of ChIPed DNA using the VAHTS Universal DNA Library Prep Kit for Illumina V3 (Vazyme Biotech, Cat. No. ND607), VAHTSTM DNA Adapters set3-set6 for Illumina (Vazyme Biotech, Cat. No. N805) and VAHTS DNA Clean Beads (Vazyme Biotech, Cat. No. N411-02) according to the manufacturer’s protocol. High-throughput sequencing was performed at Novagene (sequencing method: NovaSeq-PE150).

ChIP-seq data analysis was performed as previously described (Yu et al., 2020; Yu et al., 2021). In brief, raw data was trimmed by fastp with following parameters: “-g -q 5 -u 50 -n 15 -l 150”. The clean data was mapped to the *Arabidopsis thaliana* reference genome (TAIR10) using Bowtie2 with the default settings (Langmead and Salzberg, 2012). Only perfectly and uniquely mapped reads were used for further analysis. A summary of the number of reads for each sample is given in Supplementary Tables 3 and 4. MACS 2.0 (Feng et al., 2012) was used for peak calling with the following parameters: “gsize = 119,667,750, bw = 300, q = 0.05, nomodel, extsize = 200.” The aligned reads were converted to wiggle (wig) formats, and bigwig files were generated by bamCoverage with “-bs 10” and “-normalize using RPKM (reads per kilobase per million)” in deepTools (Ramirez et al., 2016). The data were imported into the Integrative Genomics Viewer (IGV) (Robinson et al., 2011) or Integrated Genome Browser (IGB) (Freese et al., 2016) for visualization. Only peaks that were present in both biological replicates (irreproducible discovery rate ≥ 0.05) were considered for further analysis. To annotate peaks to genes, ChIPseeker was used with default settings (Yu et al., 2015). Differential occupancy was determined using DiffBind with default settings (Ross-Innes et al., 2012). Venn diagrams were created using Venny (v.2.1) (https://bioinfogp.cnb.csic.es/tools/venny/index.html) to compare overlaps between different groups of genes. ComputeMatrix and plotProfile (Ramírez et al., 2016) were used to compare the mean occupancy density (details are shown in each corresponding figure legends).

To analyze read density and correlation between different ChIP-seq samples, we performed spearman correlation analysis. Reads density was analyzed over the merged set of binding sites across all ChIPs using multiBigwig-Summary function from deepTools. The heatmap of spearman correlation was generated by PlotCorrelation function in deepTools (Ramírez et al., 2016). Peak overlaps were analyzed by Bedtools intersect function.

### Gene ontology analysis

Gene Ontology (GO) enrichment analysis was performed using the DAVID Bioinformatics Resources (https://david.ncifcrf.gov/) and plotted at HIPLOT (hiplot-academic.com).

### Phylogenetic analysis

The amino acid sequences of orthologs proteins of SWI3D in different species were downloaded from UniProt database and used for phylogenetic analysis. The phylogenetic tree was constructed with MEGA11 (Tamura et al., 2021) using the neighbor-joining method with 1000 bootstrap replicates and the Poisson model.

### Data availability

The ChIP-seq and RNA-seq datasets have been deposited in the Gene Expression Omnibus under accession no. GSE218841 and GSE218842, respectively. The mass spectrometry proteomics data have been deposited in the Integrated Proteome Resources under the dataset identifier IPX0005495000. The BRD1, BRD2, and BRD13 ChIP-seq data and *brm-1* RNA-seq data were downloaded from GEO under accession no. GSE161595. BRIP1 and BRIP2 ChIP-seq data were downloaded from GEO under accession no. GSE142369. The H3K27me3 ChIP-seq data were downloaded from GEO under accession no. GSE145387. The H3K4me3 ChIP-seq data were downloaded from GEO under accession no. GSE183987. The H3K4me1 and H3K4me2 ChIP-seq data were downloaded from DDBJ databases under the accession number DRA010413. The H3K36me3 ChIP-seq data and *minu1-2 minu2-1* RNAseq data were downloaded from GEO under accession no. GSE205112. The H3K9me1 ChIP-seq data were downloaded from GEO under accession no. GSE146948. The H3K9ac, H3K27ac, H4K5ac and H4K8ac ChIP-seq data were downloaded from GEO under accession no. GSE183987.

### ACCESSION NUMBERS

Accession numbers of genes reported in this study include: AT2G46020 (*BRM*), AT1G21700 (*SWI3C*), AT3G01890 (*SWP73A*), AT3G03460 (*BRIP1*), AT5G17510 (*BRIP2*), AT1G20670 (*BRD1*), AT1G76380 (*BRD2*), AT5G55040 (*BRD13*), AT2G28290 (*SYD*), AT4G34430 (*SWI3D*), AT5G07940 (*SSIP1*), AT5G07970 (*SSIP2*), AT5G07980 (*SSIP3*), AT3G06010 (*MINU1*), AT5G19310 (*MINU2*), AT2G47620 (*SWI3A*), AT2G33610 (*SWI3B*), AT3G17590 (*BSH*), AT3G18380 (*SHH2*), AT1G58025 (*BRD5*), AT3G52100 (*TPF1*), AT3G08020 (*TPF2*), AT1G50620 (*OPF1*), AT3G20280 (*OPF2*), AT1G32730 (*PSA1*), AT1G06500 (*PSA2*), AT5G14170 (*SWP73B*), AT3G22990 (*LFR*), AT1G01160 (*GIF2*), AT1G18450 (*ARP4*), AT3G60830 (*ARP7*), AT4G22320 (*BCL7A*), AT5G55210 (*BCL7B*), AT3G18780 (*ACTIN2*) and AT5G09810 (*ACTIN7*).

## AUTHOR CONTRIBUTIONS

C.L. conceived the project. W.F. and Y.Y. performed most of the experiments. J.S. generated *SYD-GFP* transgenic lines. W.F., Y.Y., Y.Z., and Z.Y. conducted bioinformatics analysis. W.F., Y.Y, J.S., Z.Y., T.Z., Y.Z., Z.Z., Z.L., Y.C., C.C. and C.L. analyzed data. C.L. wrote the manuscript.

## ACKNOWLEDGEMENTS

We thank the Arabidopsis Biological Resource Center (ABRC) for seeds of T-DNA insertion lines. This work was supported by the National Natural Science Foundation of China to C.L. (32270322, 32070212, and 31870289) and to Y.Y. (32200279), the Guangdong Basic and Applied Basic Research Foundation to C.L. (2021A1515011286) and to Y.Y. (2021A1515110386), Postdoctoral Innovation Talents Support Program to Y.Y. (BX2021396), and the Fundamental Research Funds for the Central Universities to C.L. (18lgzd12).

---

**Supplemental Figure 1.**
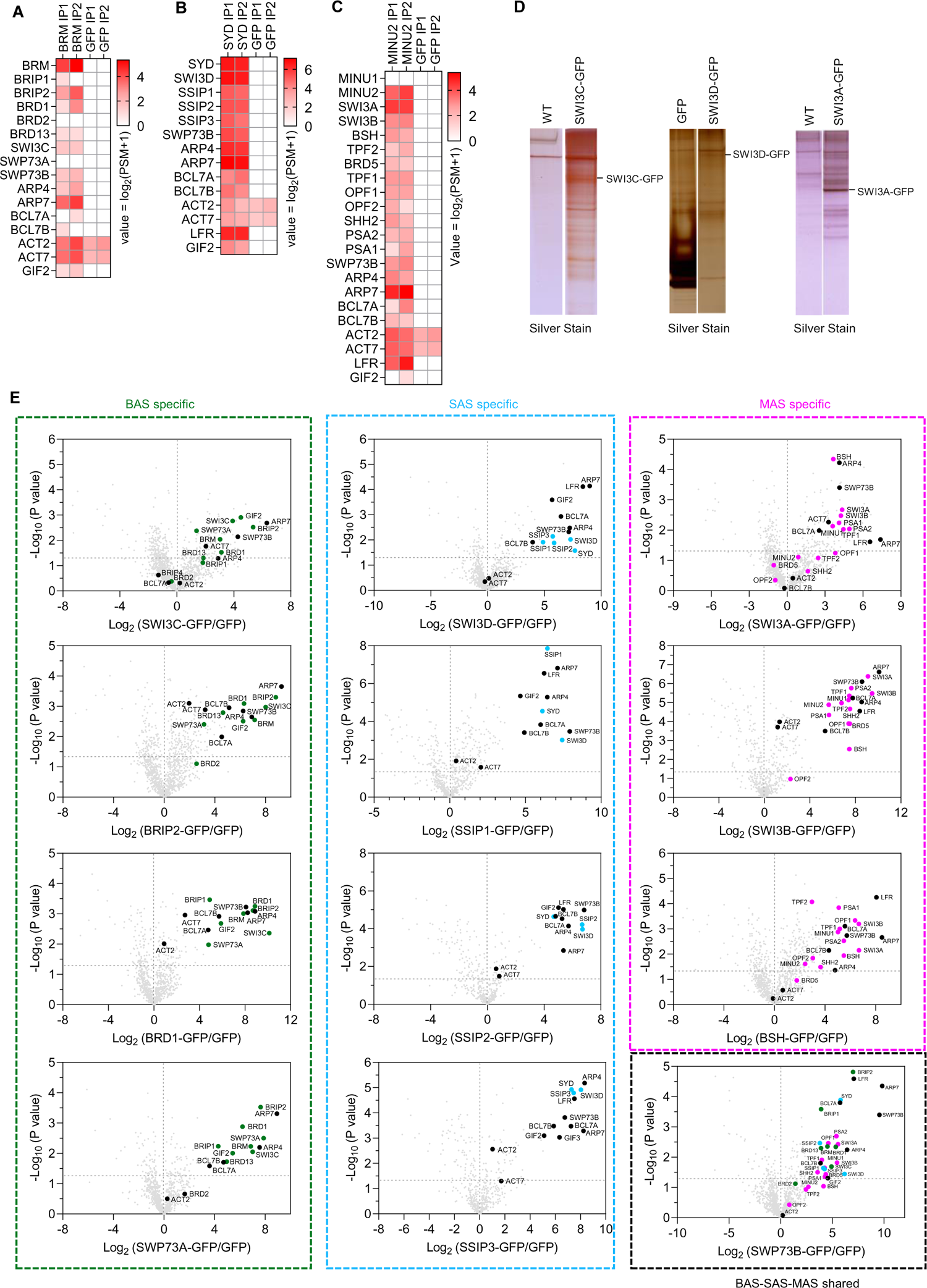
Immuno-purification of three distinct SWI/SNF sub-complexes in Arabidopsis. **A-C**, Heatmaps showing the log2(PSM+1) values of SWI/SNF complex subunits identified by IP-MS in BRM, SYD and MINU2. **D**, Sliver-stained gel of GFP immunoprecipitations from SWI3C-GFP, SWI3D-GFP and SWI3A-GFP. WT or *pACTIN2:GFP* were used as a control. **E**, Volcano plots displaying that SWI/SNF subunits are enriched in GFP immunoprecipitations from corresponding subunit from two or three independent experiments. *P* values were calculated by two-tailed Student’s t-test. The pan-SWI/SNF subunit and complex specific subunits were shown in different colors.

**Supplemental Figure 2.**
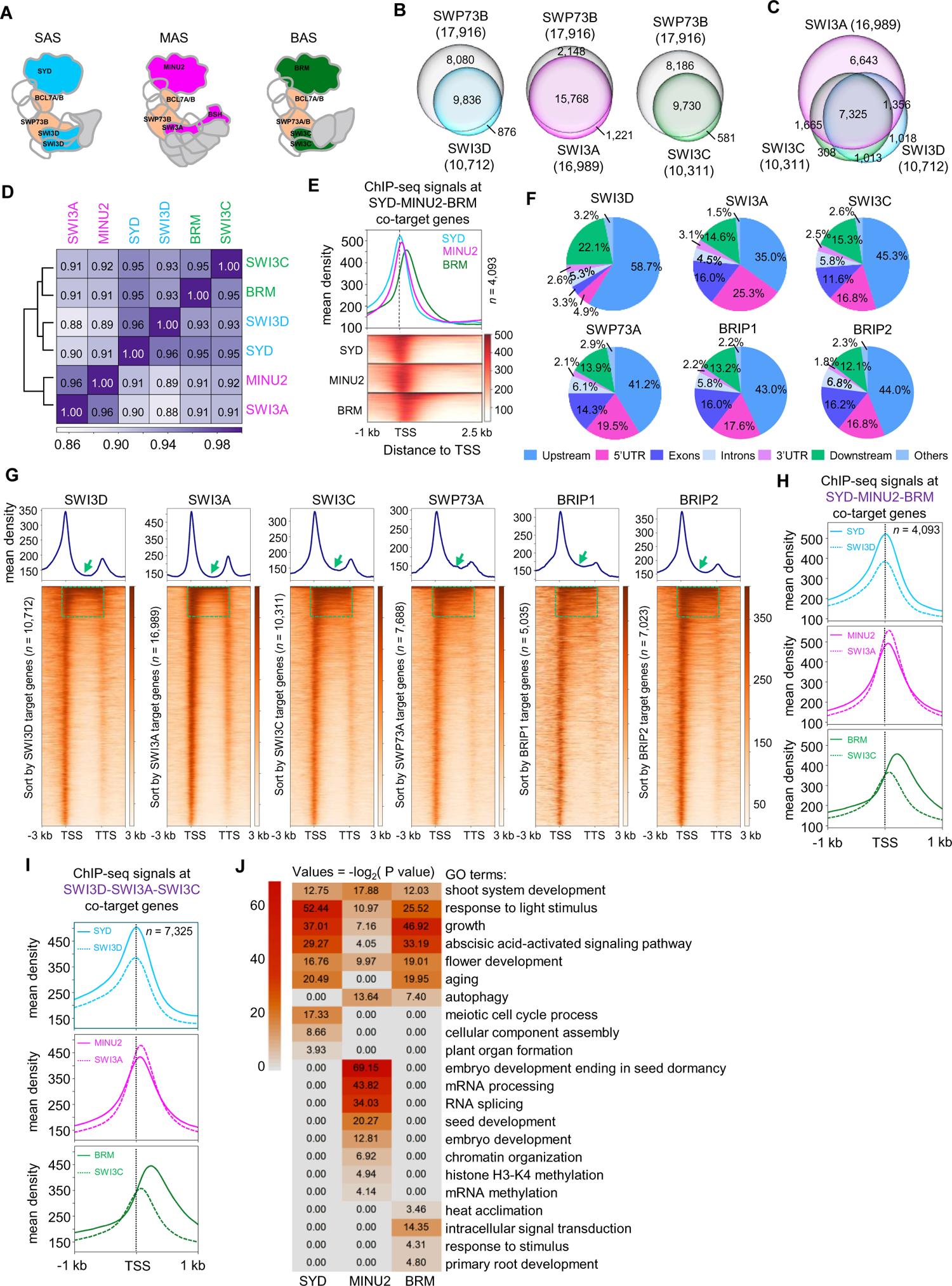
Differential binding of SAS, MAS and BAS sub-complexes on chromatin. **A**, Schematic of subunits selected for ChIP-seq: SYD and SWI3D (SAS-specific), MINU2, SWI3A and BSH (MAS-specific), BRM, SWI3C and SWP73A (BAS-specific) and SWP73B and BCL7A/B (pan-SWI/SNF) subunits. **B**, Venn diagrams displaying statistically significant overlaps among genes occupied by SWI3D, SWI3A or SWI3C with those by SWP73B. **C**, Venn diagrams displaying statistically significant overlaps among genes occupied by SWI3A, SWI3C and SWI3D. **D**, Matrix depicting spearman correlation coefficients between ChIP-seq datasets, calculated using the bin mode (bin size = 1,000). **E**, At the top, ChIP-seq read density distribution over the TSS and 2.5 kb into the gene body at their co-target genes. At the bottom, heatmap showing the mean occupancy signals of SYD, MINU2 and BRM. **F-G**, Pie charts (F), metagene plot (G) and heatmap (G) showing the distribution of SWI3A, SWI3C, SWI3D, SWP73A, BRIP1 and BRIP2 peaks at genic and intergenic regions in the genome. Green arrows and dashed boxes indicate the gene body. **H**, Metagene plot representations of the mean occupancy of SYD, SWI3D, MINU2, SWI3A, BRM and SWI3C at SYD-MINU2-BRM co-target genes. **I**, Metagene plot representations of the mean occupancy of SYD, SWI3D, MINU2, SWI3A, BRM and SWI3C at SWI3D-SWI3A-SWI3C co-target genes. **J**, Heatmap showing Gene ontology analysis of BRM, SYD and MINU2 targets genes.

**Supplemental Figure 3.**
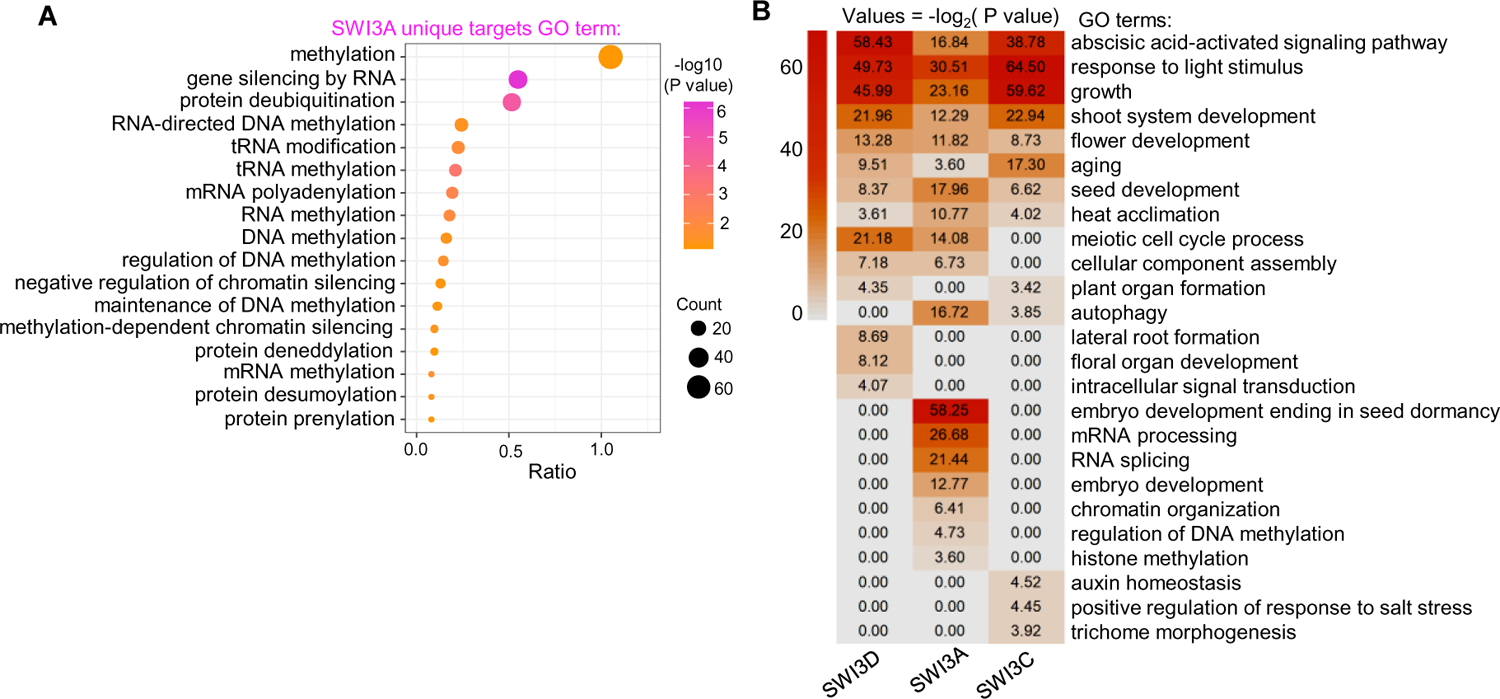
Unique target genes bound by SWI3A. **A**, Gene ontology analysis of SWI3A unique target genes. B, Gene ontology analysis of SWI3A, SWI3C and SWI3D target genes.

**Supplemental Figure 4.**
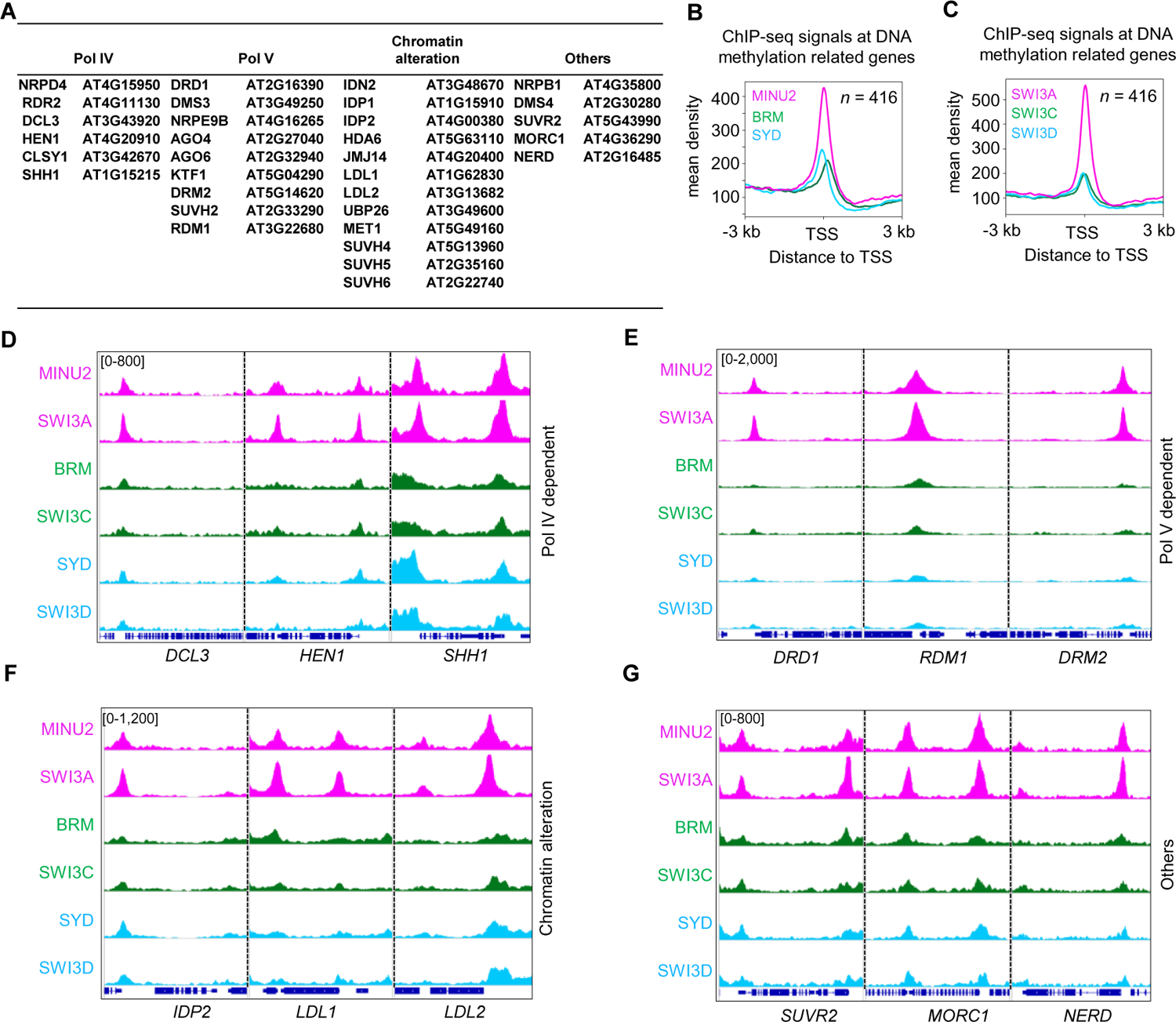
The MAS sub-complex showed a preference for binding to genes involved in the regulation of DNA methylation. **A**, Partial list of MINU2 target genes related to DNA methylation. B-C, Metagene plots displaying the mean occupancy of MINU2, BRM and SYD at the 416 genes involved in the regulation of DNA methylation. The 416 genes list were obtained by searching the GO term “Methylation” at http://geneontology.org/. D-G, IGV views of ChIP-seq signals of MINU2, SWI3A, BRM, SWI3C, SYD and SWI3D at representative genes.

**Supplemental Figure 5.**
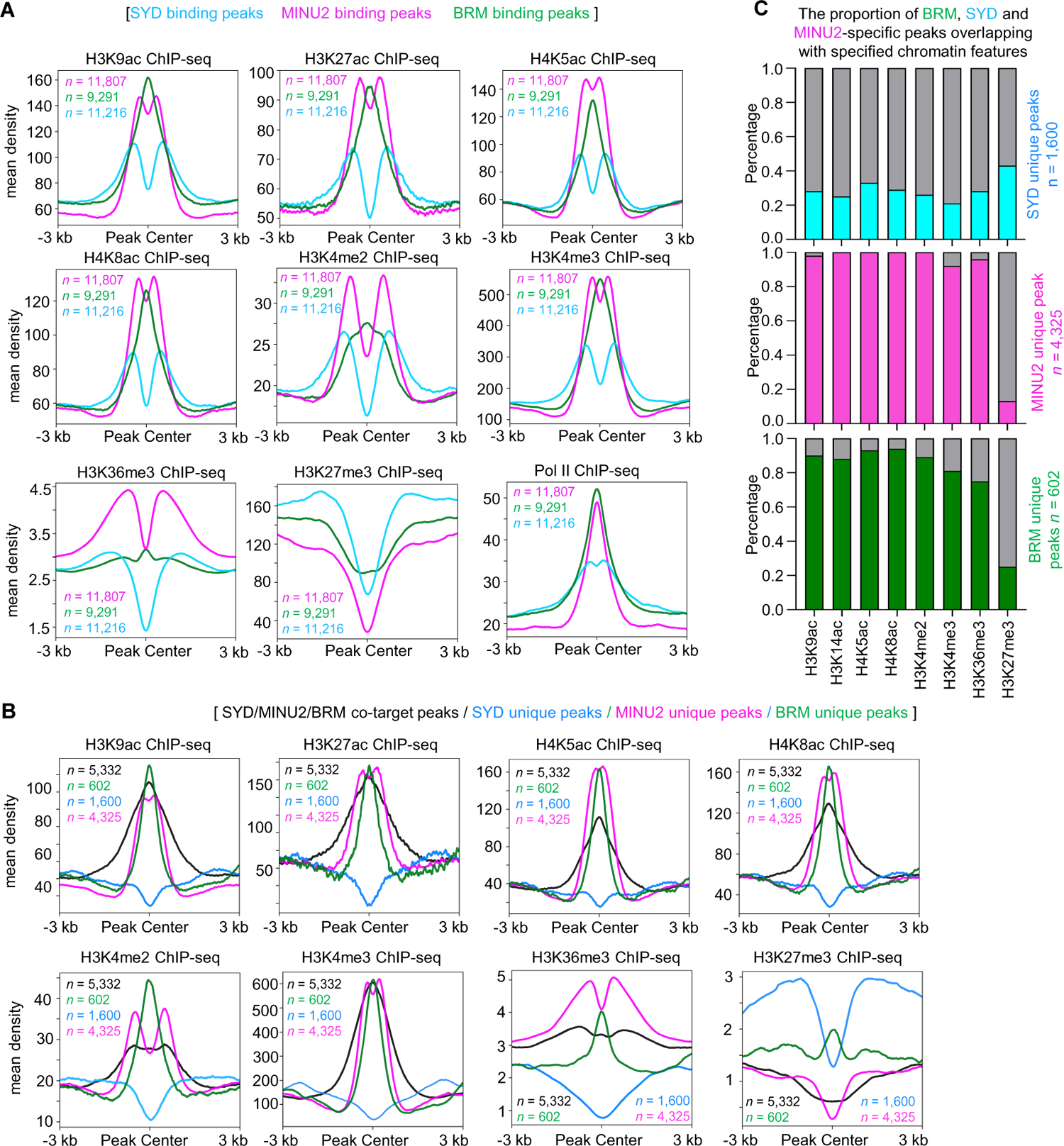
The MAS and BAS sub-complex, but not SAS sub-complex, show a significant overlap with activate histone modification markers. **A,** Metagene plots displaying the ChIP-seq signals of different histone modifications at SYD, MINU2 and BRM binding peaks. **B,** Metagene plots displaying the ChIP-seq signals of different histone modifications at SYD-MINU2-BRM co-binding and their specific binding peaks.

**Supplemental Figure 6.**
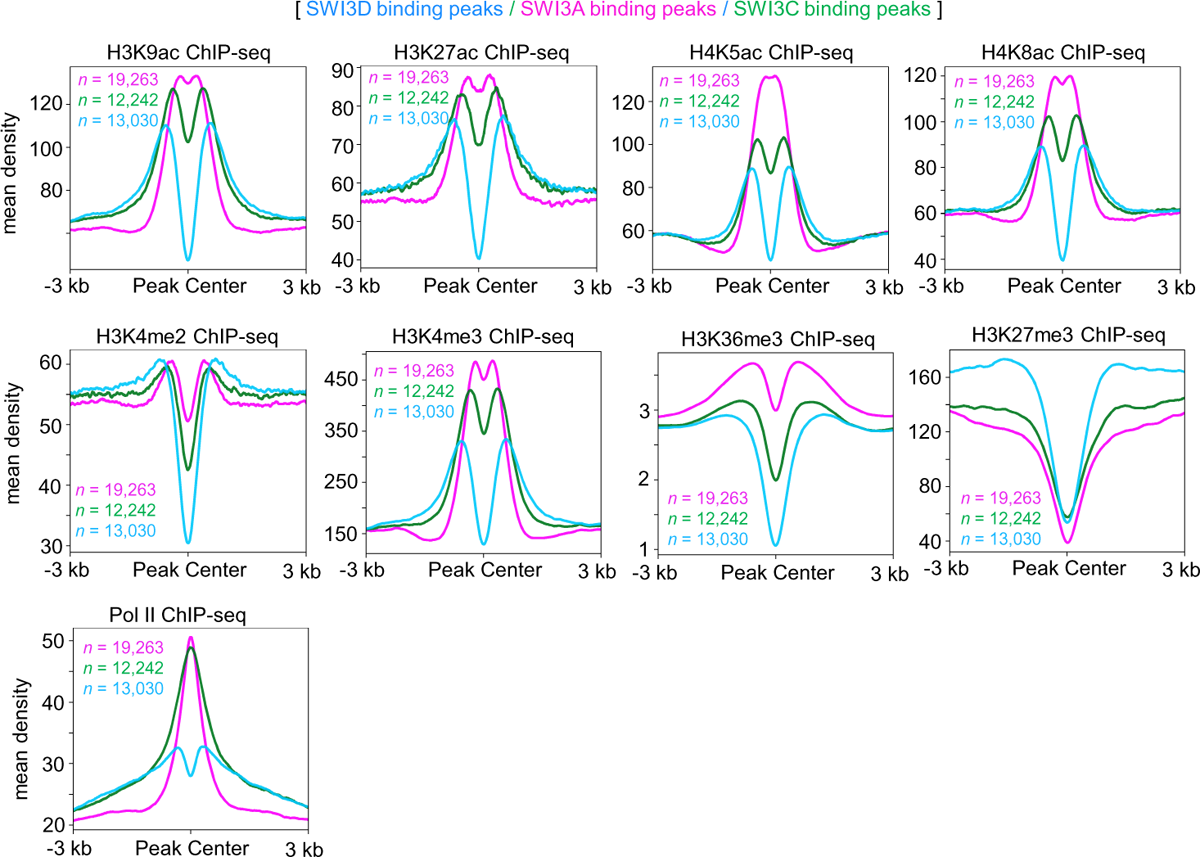
Metagene plots displaying the ChIP-seq signals of different histone modifications at SWI3D, SWI3A and SWI3C binding peaks.

**Supplemental Figure 7.**
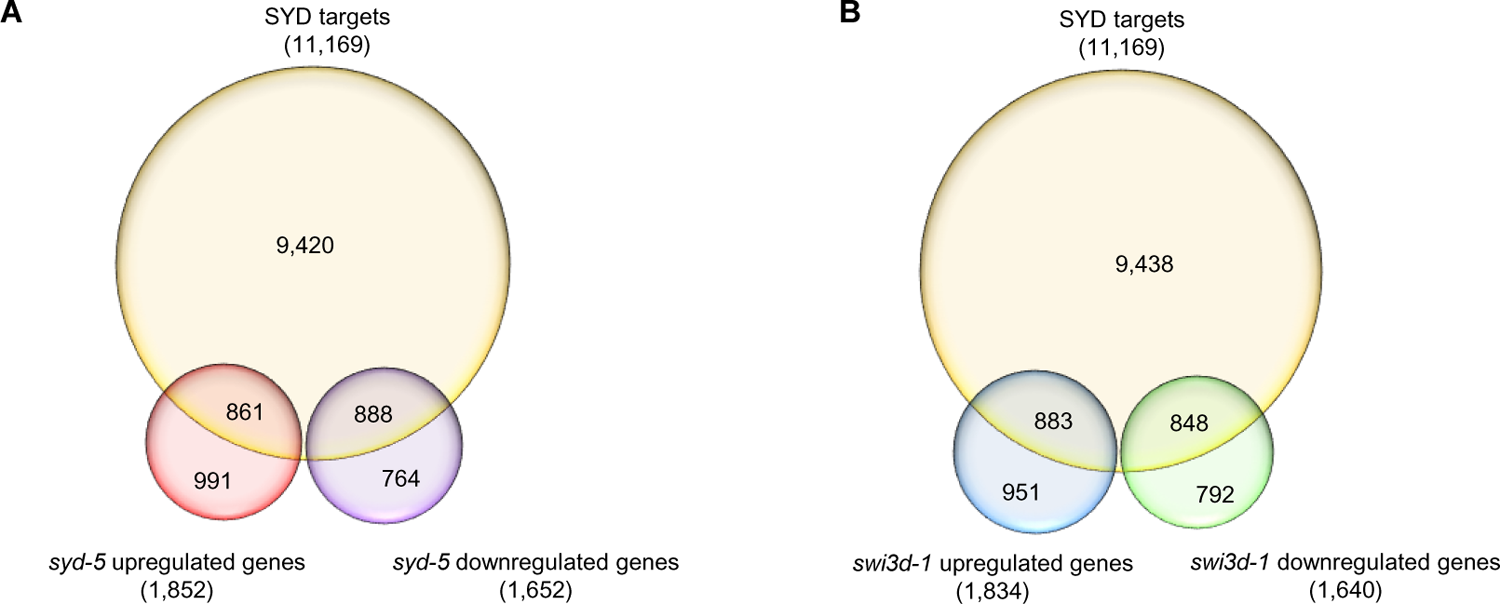
Analysis of the overlaps between SYD target genes and mis-regulated genes in *syd-5* or *swi3d-1*. **A**, Venn diagrams displaying the overlap between the genes occupied by SYD and mis-regulated genes in *syd-5*. **B**, Venn diagrams displaying the overlap between the genes occupied by SYD and mis-regulated genes in *swi3d-1*.

**Supplemental Figure 8.**
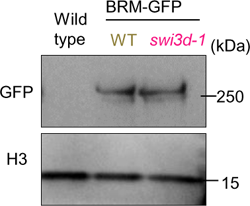
Loss of SWI3D did not affect BRM protein level. Immunoblot analysis showing the relative protein levels of BRM-GFP in a WT and *swi3d-1* background. WT was used as a GFP-free control.

**Supplemental Figure 9.**
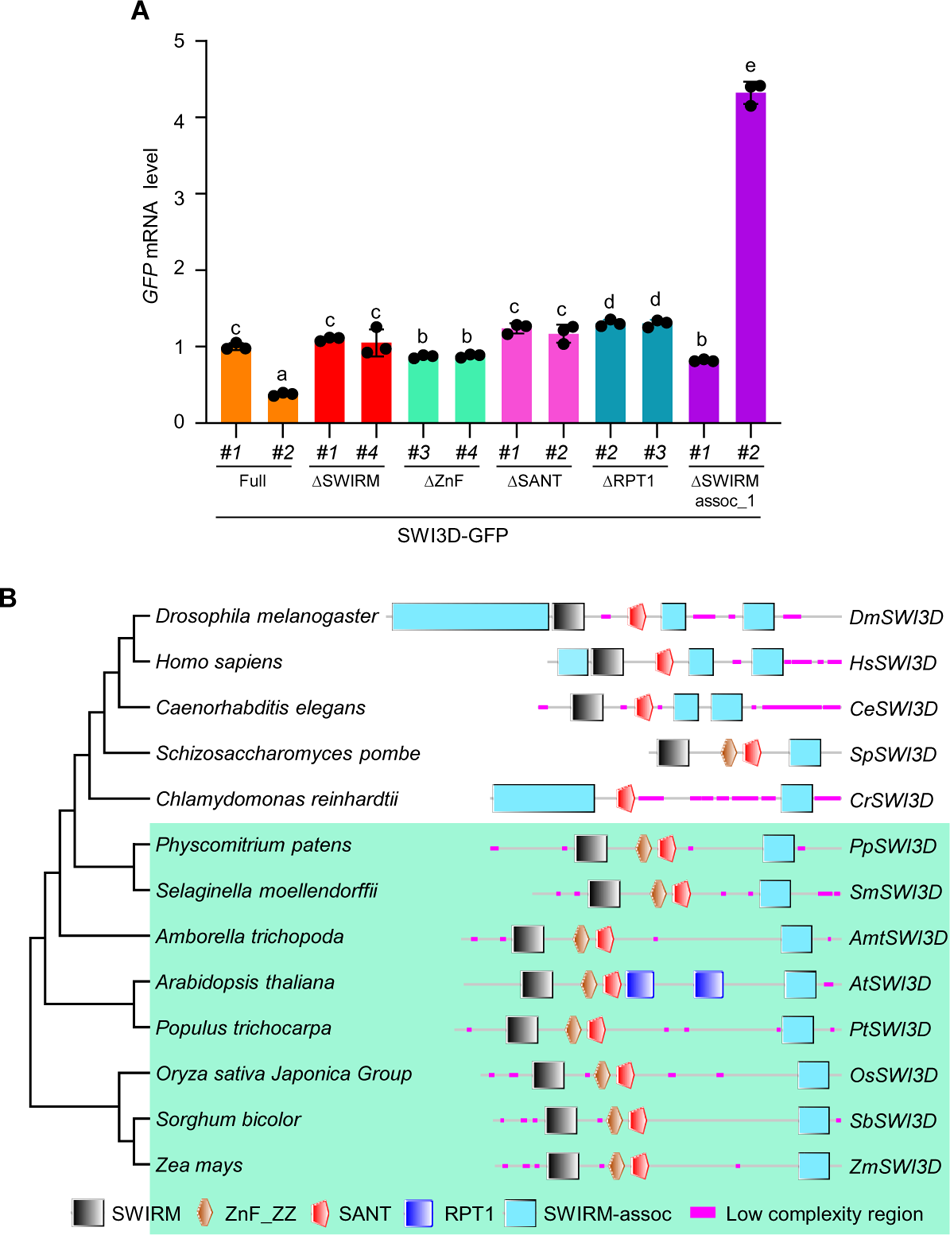
The SANT and SWIRM assoc-1 domain of SWI3D are conserved in eukaryotes. **A**, The mRNA levels of *SWI3D-GFP* were determined by RT-qPCR in different truncated versions of SWI3D. *ACTIN2* was amplified as an internal control. Error bars are presented as mean values ± s.d. from three biological replicates. Lowercase letters indicated significant differences between genetic backgrounds, as determined by the *post hoc* Tukey’s HSD test. **B**, The phylogenetic tree and genes structure of SWI3D was constructed using the amino-acid sequences from different species, including *Amborella trichopoda*, *Physcomitrella patens*, *Arabidopsis thaliana*, *Oryza sativa Japonica*, *Zea mays*, *Homo sapiens*, *Drosophila melanogaster*, *Caenorhabditis elegans*, *Schizosaccharomyces pombe*, *Chlamydomonas reinhardtii*, *Selaginella moellendorffii*, *Populus trichocarpa*, *Sorghum bicolor*. The conserved domains were predicted at the online tool SMART: https://smart.embl.de.

**Supplemental Figure 10.**
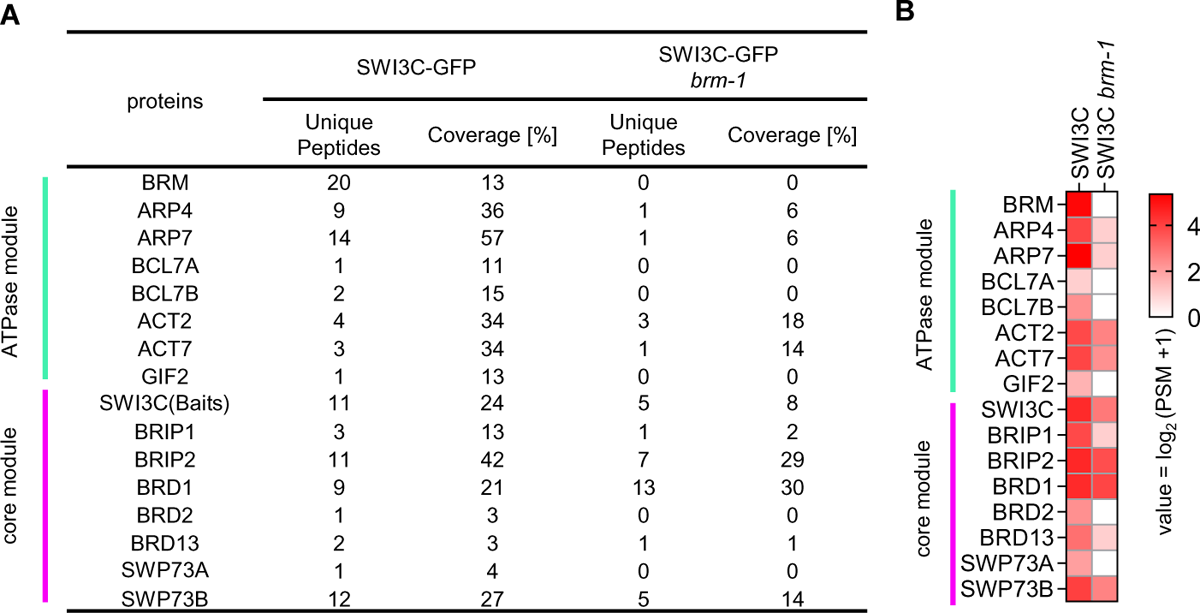
The ATPase of BAS sub-complex is essential to the stability of the core module. **A**, Unique peptide numbers of BAS sub-complex subunits identified by IP-MS in *SWI3C-GFP* under WT and *brm-1* background. **B**, Heatmap showing the log_2_(PSM+1) values of representative BAS sub-complex subunits identified by IP-MS in *SWI3C-GFP* under WT and *brm-1* background.

**Supplemental Figure 11.**
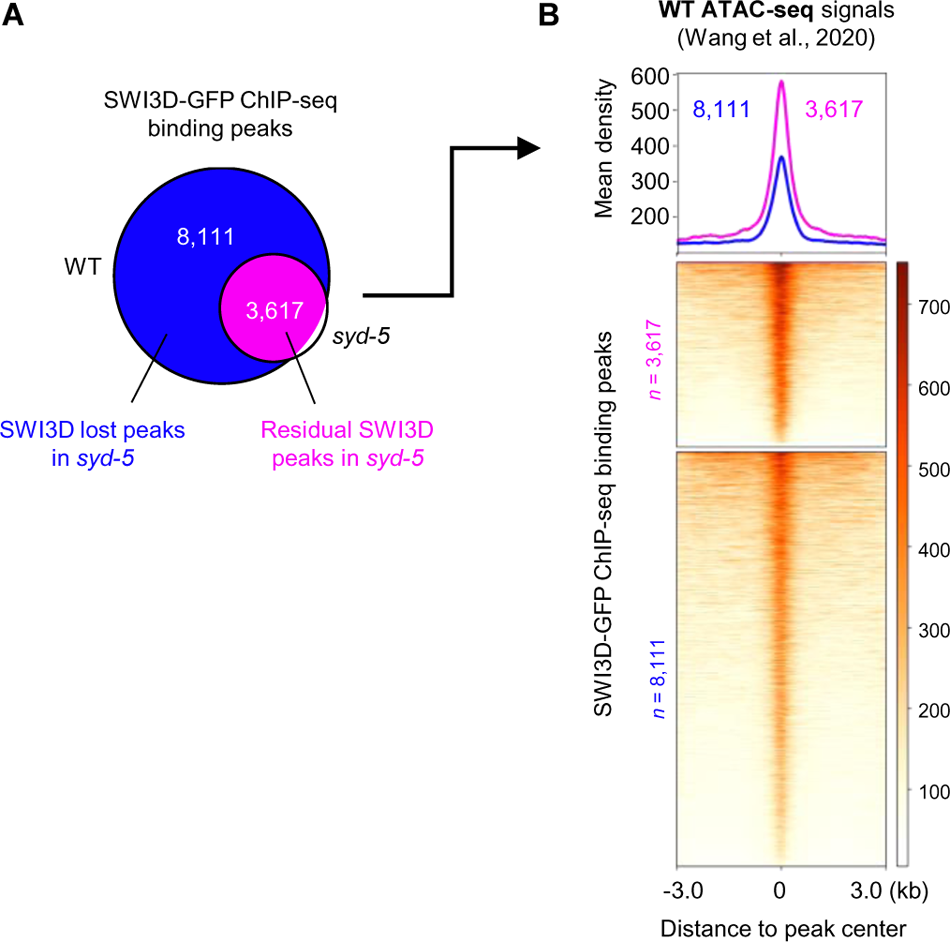
The residual peaks of SWI3D enriched at high chromatin accessibility regions. **A**, Venn diagram displaying the overlap SWI3D peaks that lost and maintained in *syd-5* mutant background. **B**, Metagene plot and heatmap representing the mean density of the ATAC-seq signals of WT at SWI3D peaks that lost and maintained in *syd-5*.

**Supplemental Figure 12.**
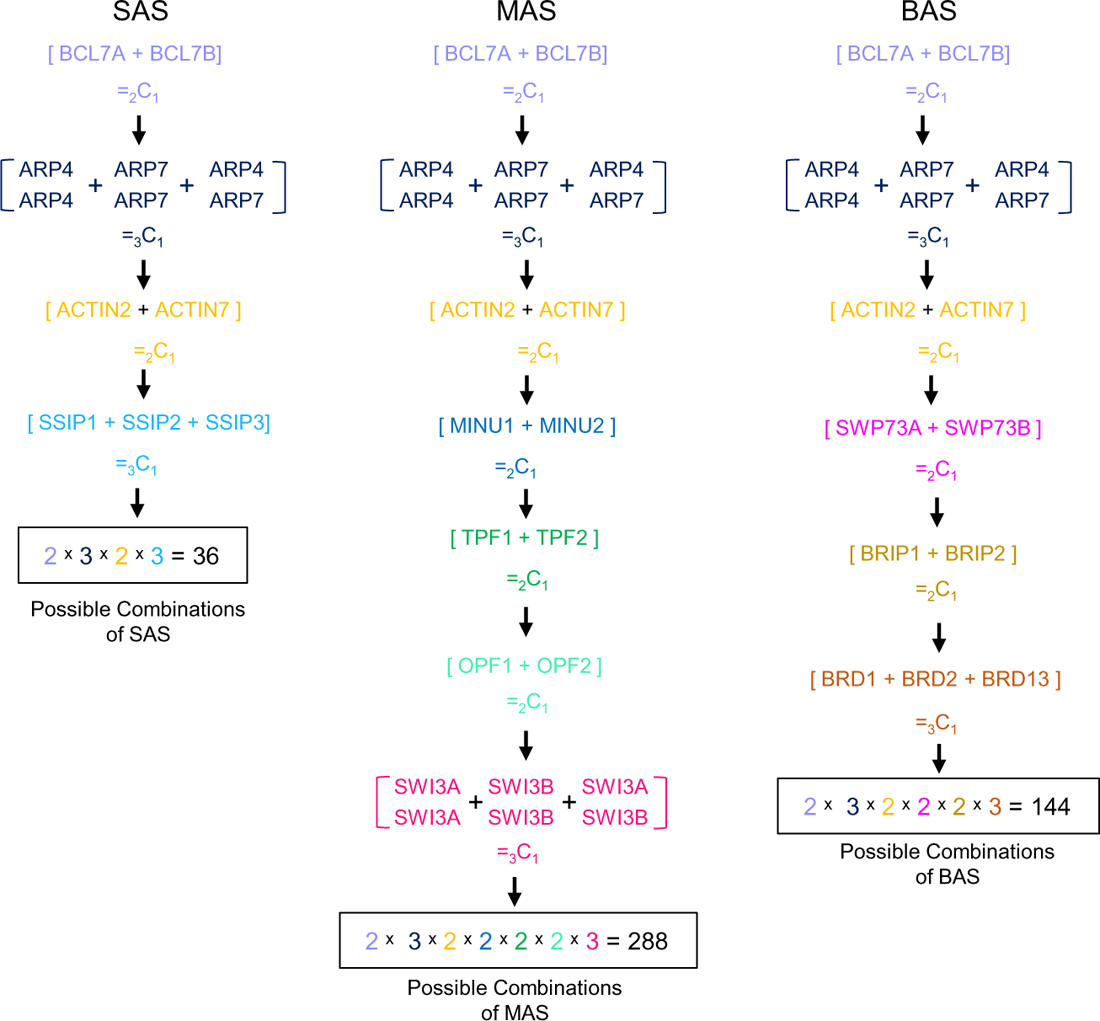
The possible combinations of the SAS, MAS and BAS sub-complexes.

**Supplemental Figure 13.**
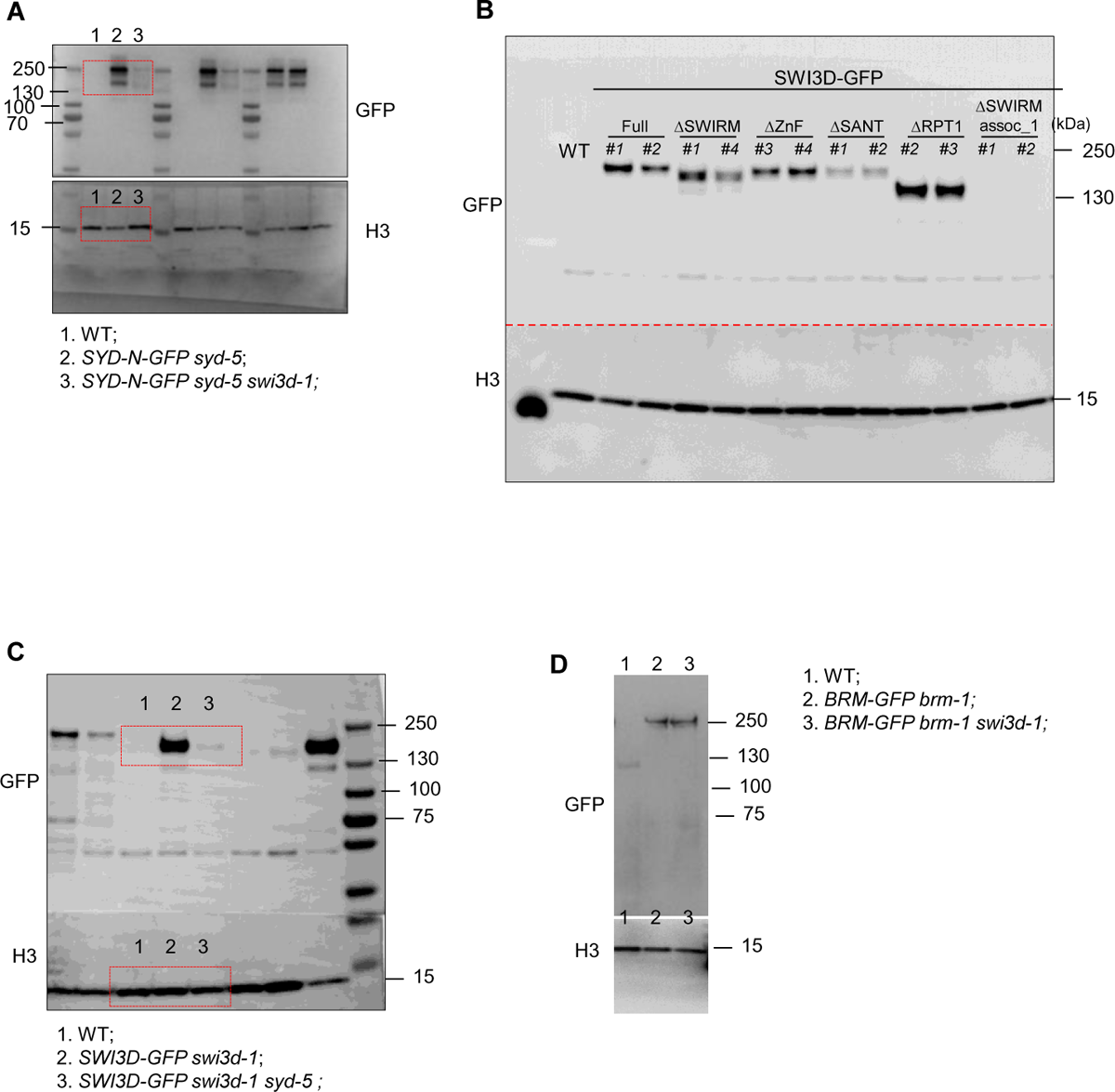
Gel source data. **A**, Western blot related to Figure 5B. B, Western blot related to Figure 6 C. **C**, Western blot related to Figure 7A. **D**, Western blot related to Supplemental Figure 8.

